# APOE interacts with COX-2 on lipid droplets to modulate inflammatory lipid signaling

**DOI:** 10.64898/2026.06.09.730852

**Authors:** Alex E. Powers, Kajal Kamble, Malvika G. Nair, Ian A. Windham, Gregory E. Miner, Courtney E. Prim, C. Allie Mills, Scott P. Lyons, Laura E. Herring, Venkat R. Chirasani, Lance A. Johnson, Sarah Cohen

## Abstract

Alzheimer’s Disease (AD) is the leading cause of dementia worldwide. Expression of the E4 variant of apolipoprotein E (*APOE*) greatly increases individuals’ risk of developing AD. In response to lipogenesis in astrocytes, APOE can escape secretion and traffic to the cytoplasmic surface of lipid droplets (LDs), but protein interactors of APOE at the LD were unknown. Here we find that LD-localized APOE physically interacts with the inflammatory lipid signaling enzyme cyclooxygenase-2 (COX-2). Like APOE, COX-2 can avoid the secretory pathway and traffic to LDs in response to lipogenesis. APOE3, but not APOE4, increases COX-2 localization to LDs. Computational modeling, microscopy-based assays, and targeted lipidomics reveal that APOE3 promotes while APOE4 suppresses COX-2 enzymatic activity at LDs and intracellular prostaglandin production. This work identifies a novel and targetable protein-protein interaction of APOE and provides a mechanistic link between *APOE4* and dysregulated inflammatory lipid signaling.

## Introduction

Alzheimer’s disease (AD) is the most common neurodegenerative disorder and the leading cause of dementia globally. AD is characterized by the progressive loss of memory and decline in cognitive function (Zheng and Wang, 2025). Over a century ago, the hallmark pathological features of AD were originally reported as extracellular amyloid-beta plaques, intraneuronal tau tangles, and accumulation of “adipose saccules” in glial cells Alzheimer Alois, 1907; Stelzmann et al., 1995). Amyloid plaques and tau tangles have been the subject of intense basic and translational investigation with mixed success in the clinic. Despite being identified at the same time, lipid accumulation has largely remained understudied. One of the leading hypotheses about the accumulation of “adipose saccules” is that this represents a pathologic increase of lipid droplets (LDs) within glia. LDs are cytoplasmic organelles that store neutral lipids including triglycerides (TGs) and cholesterol esters (CEs) within a hydrophobic core, surrounded by a phospholipid monolayer (Thiam et al., 2013; Olzmann and Carvalho, 2019; Henne and Cohen, 2026). There is an intricate protein network associated with the phospholipid monolayer of LDs that is responsible for LD turnover, cell signaling, and inflammatory signaling (Bersuker et al., 2018; Jarc and Petan, 2020a).

Recent studies have shown that both astrocytes and brain-resident immune cells, microglia, accumulate LDs with age and disease (Prakash et al., 2025; Haney et al., 2024; Wu et al., 2025; Golden et al., 2025; Marschallinger et al., 2020). Astrocytes are a glial cell type that are responsible for coordinating lipid synthesis, storage, and signaling in the brain (Ralhan et al., 2021). Astrocytes synthesize lipids, such as phospholipids and cholesterol, that are transported to neurons and used to generate new synapses, expand neuronal membranes, and maintain homeostasis (Pfrieger and Barres, 1997). Additionally, astrocytes take up lipids that have been oxidized by reactive oxygen species generated during neuronal activity to protect neurons and detoxify the lipids, preventing further damage (Nakato et al., 2015; Mauch et al., 2001; Ioannou et al., 2019; Liu et al., 2015). One of the major proteins involved in lipid homeostasis in the brain is apolipoprotein E (APOE)(Xu et al., 2006b). There are three variants of *APOE* in the human population: *APOE2, APOE3, and APOE4*. Each variant of APOE confers a different risk for developing AD (Belloy et al., 2023). *APOE3* is the most common variant and is considered a neutral risk for AD development. *APOE4* is the strongest genetic risk factor for the development of AD, with homozygous expression increasing the risk of AD up to 16-fold, while *APOE2* is protective against AD (Corder et al., 1994). The E2 and E4 variants differ from E3 by only one amino acid (APOE3 to APOE4 C112R, APOE3 to APOE2 R158C) (Weisgraber et al., 1981). It remains unclear how the *APOE4* single nucleotide variant drastically increases the risk of AD.

Astrocytes are the cell type in the brain that express the highest levels of APOE (Xu et al., 2006). Most previous work has focused on the effects of astrocyte-secreted APOE on neuronal function (Windham and Cohen, 2024; Raulin et al., 2022). However, we recently discovered that in response to conditions that stimulate lipid synthesis, APOE can escape secretion and instead traffic to the cytoplasmic surface of LDs in astrocytes (Windham et al., 2024). *APOE4* astrocytes that were pulse-chased with oleate to stimulate LD biogenesis followed by turnover displayed fewer, larger, LDs compared to APOE3-expressing cells (Windham et al., 2024). The APOE4 LDs contained higher levels of TGs, particularly TGs containing polyunsaturated fatty acids (PUFAs)(Windham et al., 2024). This increase in PUFA levels led to a higher sensitivity to lipid peroxidation in APOE4-relative to APOE3-expressing astrocytes, suggesting a metabolic rewiring of astrocytes by APOE4 that results in higher lipid load and more lipid peroxidation (Windham et al., 2024). In addition, lipidomic and proteomic studies revealed dramatic differences in the lipidome and proteome between APOE3- versus APOE4-expressing LDs isolated from liver or astrocytes, respectively (Friday et al., 2025; Cuní-López et al., 2025). However, the mechanism by which LD-localized APOE modulates droplet lipid composition is unknown.

In this study, we sought to understand how LD-associated APOE modulates LD composition. To identify binding partners of APOE on LDs, we performed affinity-purification of APOE followed by proteomics under conditions where APOE localized to LDs. We identified the inflammatory lipid signaling enzyme, prostaglandin-endoperoxide synthase-2 (PTGS2) – more commonly known as cyclooxygenase-2 (COX-2) – as the top binding partner of both APOE3 and APOE4 at the LD. Using BioEmu, we modeled the APOE-COX-2 interaction on a lipid interface and observed differential binding characteristics between APOE3 and APOE4.

Immunocytochemistry revealed that APOE3 increased COX-2 localization to LDs, while APOE4 did not. Consistent with this result, targeted lipidomics for arachidonic acid metabolites revealed that APOE3 increased the intracellular production of signaling lipids downstream of COX-2, relative to APOE4. Our study contributes to the growing body of literature that lipid dysfunction in astrocytes contributes to AD pathology and provides a novel mechanism by which APOE4 contributes to metabolic rewiring and dysregulated inflammation.

## Results and Discussion

### APOE interacts with COX-2 at lipid droplets

APOE can reroute from the secretory pathway to LDs in states of lipogenesis, such as exogenous fatty acid treatment (Windham et al., 2024). We previously discovered that APOE can influence the size and composition of LDs in astrocytes, with APOE4 increasing the polyunsaturated fatty acid composition of TGs relative to APOE3 (Windham et al 2024). However, the mechanism by which APOE influences LD composition remains unclear. We hypothesized that APOE modulates LD lipid composition through interaction with an enzyme on the LD surface. To identify candidate binding partners of APOE at the LD, we performed affinity-purification proteomics. As our model system, we used APOE targeted replacement astrocytes, which express human APOE3 (TRAE3) or APOE4 (TRAE4) at the mouse endogenous locus (Morikawa et al., 2005). We virally transduced TRAE3 cells to express either mEmerald (Em) or mEmerald-tagged APOE3 (APOE3-Em), while TRAE4 cells were transduced with Em or mEmerald-tagged APOE4 (APOE4-Em)(Fig. 1, A). Transduced cells were then incubated with the common dietary fatty acid oleic acid (OA) to drive APOE onto LDs, or without OA as a control. APOE3-Em, APOE4-Em, or mEmerald was immunoprecipitated, and putative binding partners identified by LC-MS/MS (Supplemental Table 1). Figure 1B summarizes the results of two independent affinity purification experiments using two different LC-MS/MS instruments. To compare across experiments, we calculated the Log2 fold-enrichment observed with OA treatment compared to untreated control cells from each experiment and averaged the values.

**Figure 1.**
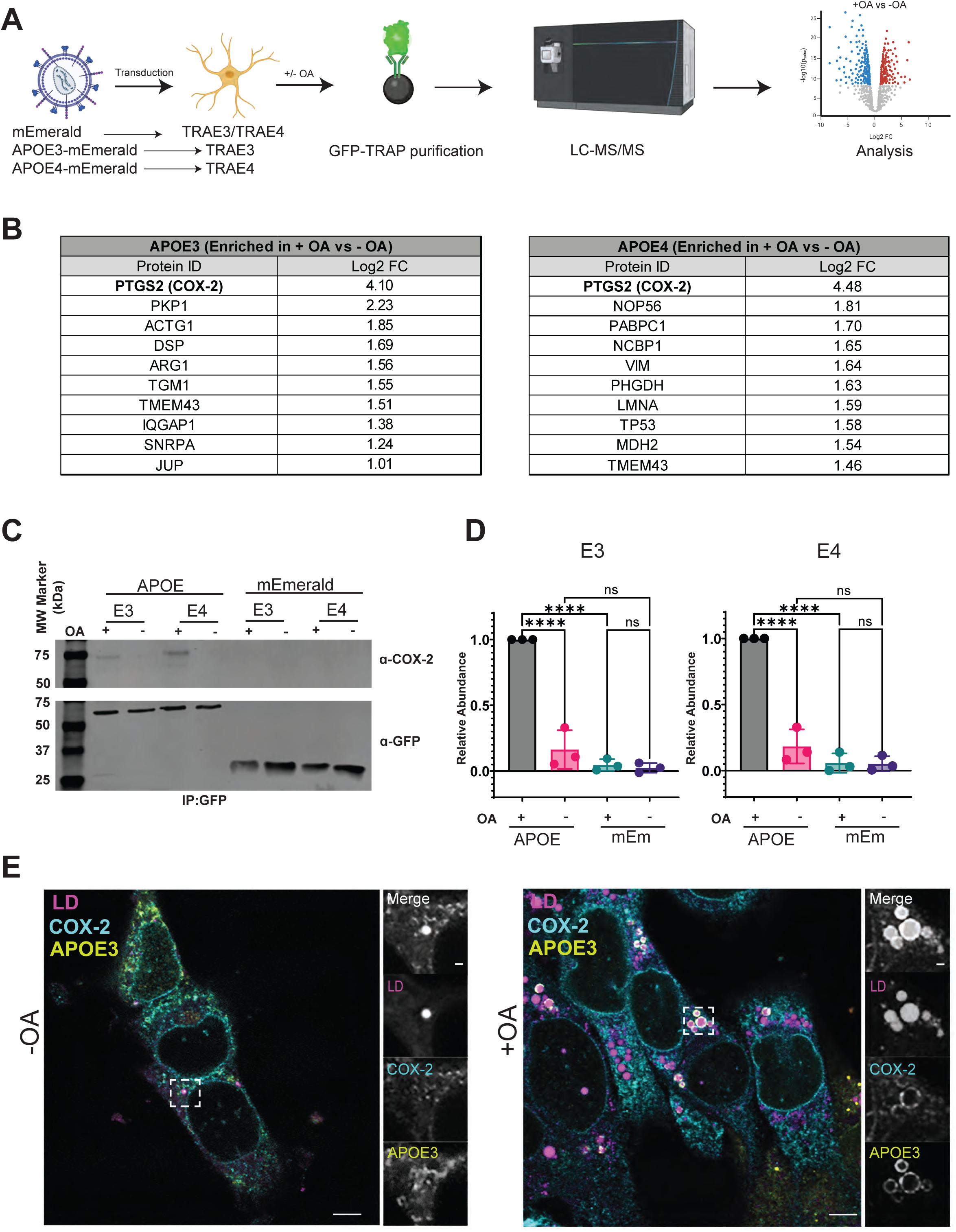
APOE interacts with COX-2 on lipid droplets. (A) Schematic of experimental workflow to identify LD-associated APOE binding partners. (B) Table of 10 most significantly enriched proteins with APOE3-mEmerald or APOE4-mEmerald after oleate pulse relative to baseline after filtering proteins with a Log2 fold change >1.0 compared to mEmerald controls. Values are the average of 2 biological replicates. (C) Representative immunoblot of GFP (mEmerald) and COX-2 detected following immunoprecipitation of APOE-mEmerald or mEmerald. (D) Quantification of intensity of COX-2 immunoprecipitation in 3 independent biological replicates, normalized to total mEmerald pulldown. (E) Representative single plane confocal images of TRAE3 cells at baseline or after 400 µM oleate pulse, immunostained for APOE and COX-2 and dyed with BODIPY493 to label LDs. Scale bar: 10µm. Groups were compared by one-way ANOVA with a post-hoc Tukey’s t test in Graphpad Prism. p ≤0.05*, p ≤0.005**, p ≤0.0005***, p ≤0.0001**** ≤0.0001****

To our knowledge, this is the first reported co-immunoprecipitation of APOE in astrocytes. Interestingly, in TRAE3 cells at baseline, some of the most highly enriched putative interacting proteins (relative to mEm control) were involved in cytoskeleton and basement membrane organization (TRIO, KIRT1, HSPG2), the ubiquitination pathway (RNF31, TRIM32), and secretion (SEC16a)(Table 1). In TRAE4 cells at baseline, as previously reported in both liver cells and tissue (Rueter et al., 2023),the most highly enriched interactors were mitochondrial associated voltage-gated anion channel proteins (VDAC1-3). Among other high confidence interacting partners of APOE4 in astrocytes at baseline were RNA splicing and ribosomal proteins (THRAP3, RPS26, RPL4, RPL14), suggesting APOE could influence gene expression at all molecular levels in these cells. With the exception of KIRT1, all these proteins were identified as putative interactors of both APOE3 and APOE4, but were enriched to varying degrees depending on APOE genotype.

To distinguish APOE function on the LD from functions in the secretory pathway, we focused our analysis on identifying binding partners enriched in the oleate treated conditions relative to the untreated conditions (Fig. 1B). Interactors of LD-associated APOE again included RNA splicing and ribosomal proteins (APOE3: SNRNPA; APOE4: NOP56, PABPC1, NCBP1). LD-associated APOE3 preferentially interacted with desmosomal proteins (PKP1, DSP, JUP), while LD-associated APOE4 bound intermediate filament proteins (VIM, LMNA). Only two of the top ten candidate proteins were shared between APOE3 and APOE4: TMEM43 and COX-2. TMEM43 is an ER transmembrane and nuclear envelope structural protein (Ratnavadivel et al., 2026), while COX-2 is a cyclooxygenase that converts arachidonic acid to prostaglandin GH2 (PGH2), which can then be funneled into additional prostaglandin inflammatory signaling lipids including PGD2, PGE2, PGF2a and PGI2 (Wang et al., 2021). Because of the rich literature implicating dysregulated inflammation in AD (Heneka et al., 2025; Lee et al., 2025), we were excited to follow up on this candidate – which also happened to be the top hit for both APOE3 and APOE4. We confirmed the APOE-COX-2 interaction by performing co-immunoprecipitation of APOE followed by detection of COX-2 by western blotting. We found that COX-2 did not co-immunoprecipitate with Em, indicating specificity of the interaction, and only co-immunoprecipitated with APOE3-Em and APOE4-Em under oleate treated conditions, supporting the idea that these proteins interact on the LD (Fig. 1, C and D).

Like APOE, COX-2 has a signal sequence and usually translocates into the ER (Michael Garavito et al., 2002). However, COX-2 has been observed on LDs in other cell types such as cancer cells and leukocytes (Accioly et al., 2008; Bozza et al., 1996). To test whether COX-2 can localize to LDs in astrocytes, we performed immunofluorescence to detect endogenous APOE and COX-2 localization in TRAE3 and TRAE4 cells. We found that like APOE, COX-2 partially relocalizes from the secretory pathway to LDs in oleate treated cells, forming characteristic LD-associated ring structures, and co-localizing with APOE (Fig. 1, E). Together, these results suggest that under lipogenic conditions, both APOE and COX-2 traffic to LDs, where they interact at the LD surface.

### Conformational landscapes of APOE3 and APOE4 are thermodynamically distinct

To elucidate the structural basis of APOE-COX-2 interactions, we first characterized the equilibrium conformational ensembles of APOE3 and APOE4 using the generative machine learning framework BioEmu. This approach is particularly appropriate for comparing APOE isoforms, because the functional distinction between APOE3 and APOE4 is not attributable to a gross change in the static crystal structure – which differs only subtly – but rather to isoform-specific differences in the dynamic conformational ensemble sampled in solution (Dong et al., 1994). Projection of the APOE3 and APOE4 conformational ensembles onto their respective two-dimensional free energy landscapes revealed thermodynamically distinct preferences between the two variants (Fig. 2, A and B). The APOE3 landscape is characterized by a well-defined global free energy minimum corresponding to the canonical elongated four-helix bundle conformation. The representative centroid structure extracted from this minimum exhibits Cys112 oriented inward, participating in hydrophobic packing interactions within the helix bundle interior that stabilize the N-terminal domain in an open, extended conformation (Fig. 2 A). This structural state is consistent with the established topology of the lipid-free APOE3 N-terminal domain, in which the four-helix bundle and the C-terminal domain do not engage in stable interdomain contacts.

**Figure 2.**
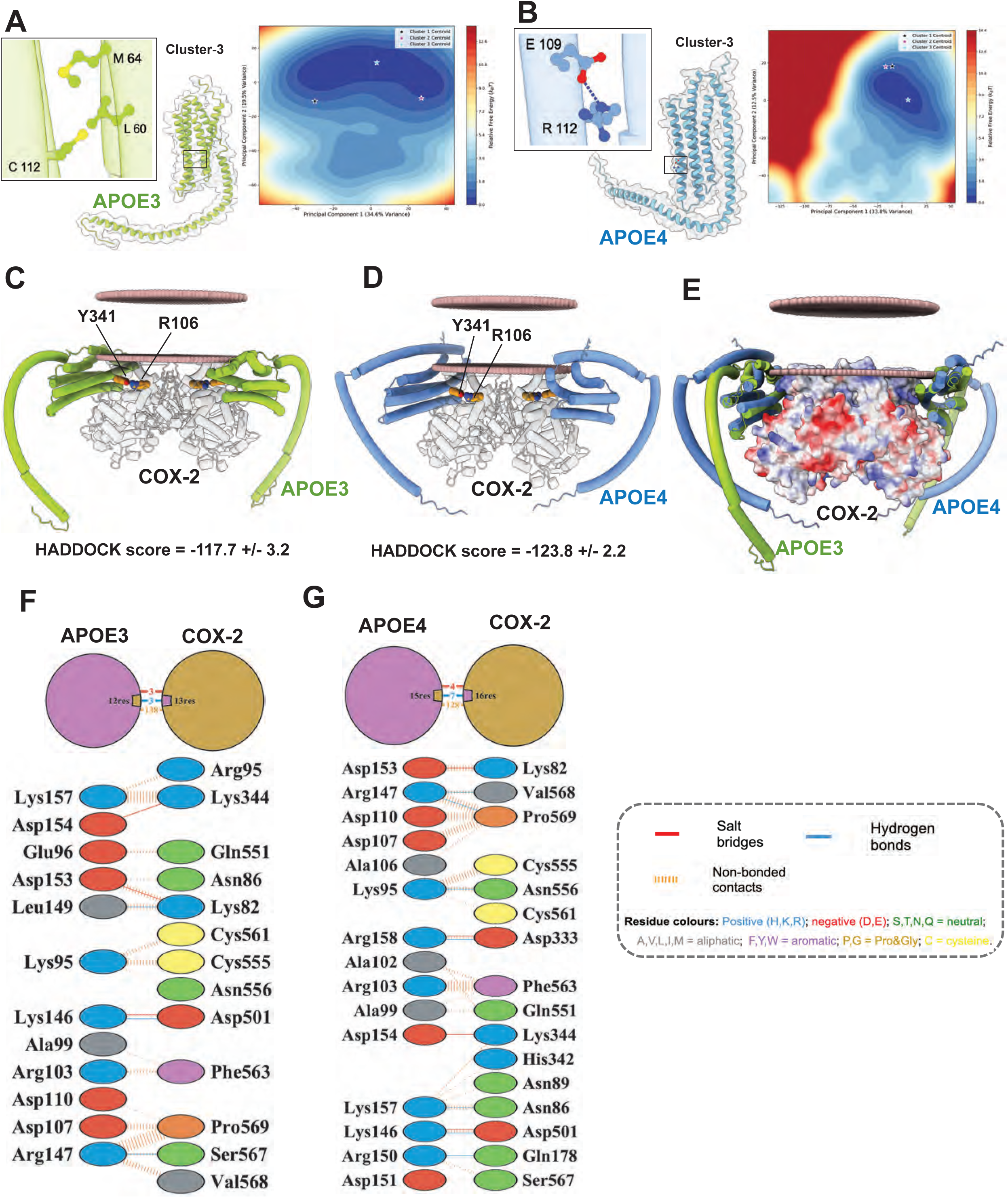
Structural and energetic characterization of APOE3 and APOE4 binding to the COX-2 homodimer. (A) BioEmu-derived free energy landscape of APOE3 projected onto the first two principal components (PC1, PC2) of the conformational ensemble. Each point represents one sampled conformation; color encodes relative free energy (ΔG, kcal/mol) computed from the Gaussian kernel density estimate of the population probability. The inset ribbon diagram shows the representative centroid structure extracted from the global free energy minimum, with Cys112 depicted in stick representation. The inward orientation of Cys112 and its participation in hydrophobic packing with Leu60 and Met64 within the four-helix bundle is indicated. (B) Free energy landscape of APOE4 constructed as in (A). The representative centroid structure illustrates the conformational consequences of the C112R polymorphism: surface exposure of Arg112, formation of a new Arg112–Glu109 salt bridge, and the resulting remodeling of the four-helix bundle geometry. The altered conformational basin relative to APOE3 reflects the isoform-specific shift in the equilibrium ensemble driven by this mutation. HADDOCK docked models of (C) APOE3 and (D) APOE4 molecules bound to the COX-2 homodimer. The COX-2 active site residues – Arg106 and Tyr341 are shown in orange spheres. The lipid bilayer boundary is approximated by a layer of spherical pseudo-atoms (brown spheres), illustrating the spatial relationship of the complex to the membrane plane. (E) Structural superposition of the APOE3–COX-2 and APOE4–COX-2 docked complexes. The COX-2 surface is colored by electrostatic potential calculated using APBS. PDBsum two-dimensional interaction diagram for the interfaces of (F) APOE3–COX-2 and (G) APOE4–COX-2.

By contrast, the APOE4 free energy landscape is dominated by a conformationally shifted global minimum driven by the C112R polymorphism. The C112R substitution introduces a guanidinium side chain at position 112 that forms a new intramolecular salt bridge with Glu109, disrupting the native hydrophobic packing of the four-helix bundle and triggering a cascade of long-range structural rearrangements (Nemergut et al., 2023). Consistent with this mechanism, the representative APOE4 centroid structure demonstrates surface exposure of Arg112 and rearrangement of the adjacent helical geometry (Fig. 2 B). This “domino-like” propagation of the C112R perturbation Nemergut et al., 2023) repositions Arg61, which in APOE3 participates in an intra-bundle Arg61-Glu109 interaction; in APOE4, the loss of this contact allows Arg61 to engage Glu255 in the C-terminal domain, establishing an aberrant interdomain salt bridge that compacts the overall molecular architecture (Nguyen et al., 2014). The resulting APOE4 ensemble thus samples a more restricted set of conformational states with an altered surface topology relative to APOE3, providing a structural basis for the differential protein–protein interaction behavior reported for the two isoforms.

### APOE3 and APOE4 engage the COX-2 homodimer through slightly altered binding modes

To investigate the structural mechanism by which APOE isoforms might differentially interact with COX-2, we docked the BioEmu-derived representative structures of APOE3 or APOE4 against the COX-2 homodimer using the HADDOCK integrative docking platform (Dominguez et al., 2003) in a 2:1 (APOE monomer:COX-2 dimer) stoichiometry. Both APOE3 and APOE4 yielded convergent, well-clustered docking solutions; however, the two isoforms exhibited divergent binding orientations and affinities at the COX-2 surface (Fig. 2, C and D). The HADDOCK score for the APOE3:COX-2 complex is -117.7 +/- 3.2, while for the APOE4:COX-2 complex is -123.8 +/- 2.2. Structural superposition of the top-ranked APOE3–COX-2 and APOE4–COX-2 complexes reveals that the two APOE variants engage COX-2 in slightly altered poses relative to the membrane plane (Fig. 2, E). Detailed characterization of the APOE–COX-2 binding interfaces using PDBsum (Laskowski et al., 2018) revealed quantitative differences in the intermolecular contact networks formed by the two isoforms (Fig. 2, F and G).

### APOE3 promotes COX-2 localization to LDs

Our structural predictions suggested the hypotheses that APOE3 and APOE4 may differentially affect COX-2 recruitment or stabilization at the LD, and/or COX-2 enzymatic activity. We first tested the hypothesis that APOE is required to recruit COX-2 to LDs. The N-terminal region of APOE shares remarkable structural similarities to perilipin-3 (PLIN3)(Faustino et al., 2015; Hickenbottom et al., 2004). PLIN3 is a resident LD protein that influences LD turnover by interacting with adipose triglyceride lipase and co-activator CGI-58 to initiate lipolysis in skeletal muscle (MacPherson et al., 2013). We reasoned that APOE could perform a similar role to PLIN3 for COX-2 in astrocytes. To test this, we knocked down endogenous APOE expression in TRAE3 or TRAE4 astrocytes via siRNA, then treated with oleate and stained for APOE, COX-2, and LDs. As expected, in both TRAE3 and TRAE4 astrocytes we observed APOE enrichment around LDs only in oleate-treated cells following transfection with non-targeting siRNA (NT-siRNA)(Fig. 3, A-B). Following APOE-siRNA-mediated knockdown, we first confirmed loss of APOE expression via loss of immunofluorescent APOE signal relative to NT-siRNA (Fig. 3, A-B). We then quantified COX-2 intensity around LDs. We found that COX-2 localization to LDs was markedly reduced in NT-siRNA-treated TRAE4 cells relative to TRAE3 cells (Fig. 3, C). Following APOE knockdown, we observed a drastic reduction in the levels of COX-2 associated with LDs in TRAE3 astrocytes; in contrast we observed no effect of APOE KD on COX-2 LD localization in TRAE4 cells (Fig. 3, C). Intriguingly, COX-2 localization in TRAE3 cells following APOE knockdown was similar to levels observed in TRAE4 cells regardless of APOE4 expression (Fig. 3, C), suggesting that APOE3 specifically can influence either the recruitment or retention of COX-2 to LDs. Total COX-2 intracellular integrated intensity was modestly increased by oleate pulse and was higher in TRAE3 cells than in TRAE4 cells under oleate-treated conditions (Fig. 3, D). This suggests that APOE can also modulate cell-wide regulation of COX-2 levels by influencing COX-2 turnover, with APOE3 stabilizing COX-2 relative to APOE4.

**Figure 3.**
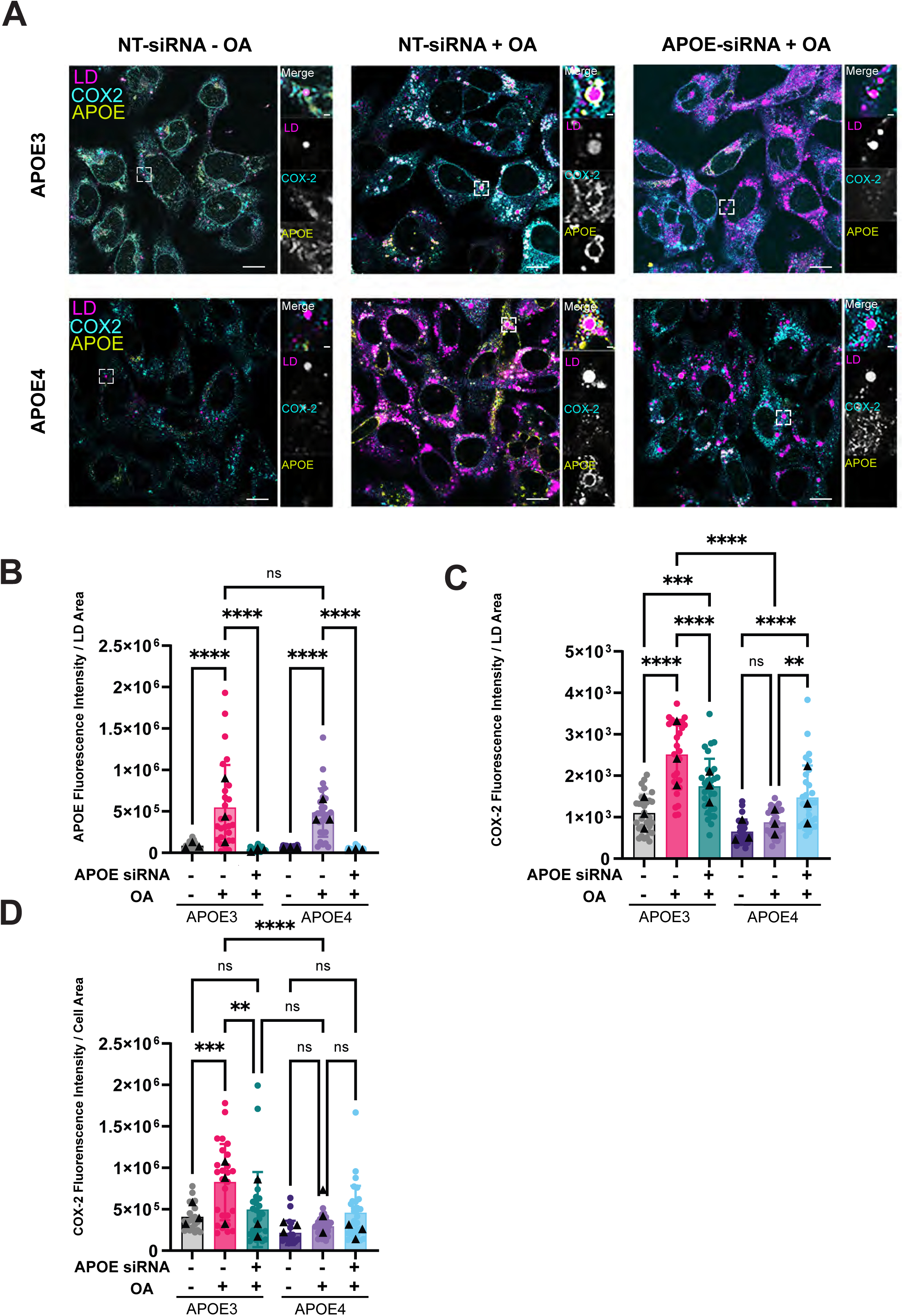
APOE3, but not APOE4, promotes COX-2 localization on lipid droplets. (A) Single plane confocal micrographs of fixed TRAE3 (top) or TRAE4 (bottom) astrocytes stained for LDs (pseudocolored magenta, BODIPY493), COX-2 (pseudocolored turquoise, Alexafluor568), and APOE (pseudocolored yellow, Alexafluor647). Left: TRAE cells at baseline after treatment with a non-targeting (NT) siRNA. Center: TRAE cells after NT siRNA treatment and a 400 µM oleate pulse for 5 hours. Right: TRAE cells after APOE siRNA knockdown and 400 µM oleate pulse for 5 hours. (B,C) Quantification of integrated intensity (arbitrary units, AU) of LD associated APOE (B) or COX-2 (C) normalized to LD area in TRAE cells by ImageJ software. (D) Quantification of integrated intensity (arbitrary units, AU) of total COX-2 normalized to cell area. Scale bars: 10 µm, 2.5 µm. Triangle: Mean of independent biological replicate. Circle: Individual cell measurement across biological replicates. Groups were compared by one-way ANOVA with a post-hoc Tukey’s t test in Graphpad Prism. p ≤0.05*, p ≤0.005**, p ≤0.0005***, p ≤0.0001****

### APOE4 phenocopies COX-2 inhibition and increases arachidonic acid containing TGs

LDs are signaling hubs that provide the fatty acid precursors for inflammatory lipid mediators (Jarc and Petan, 2020b). Since we observed a significant reduction in COX-2 recruitment to LDs in TRAE4 astrocytes, we hypothesized that this could result in lower COX-2 activity at LDs. We previously reported that expression of APOE4 causes larger LDs and increased total LD area upon oleate pulse-chase, particularly after the oleate chase (Windham et al., 2024). Therefore, we next investigated whether pharmacological inhibition of COX-2 via aspirin phenocopied the APOE4 LD phenotype. We treated APOE3 or APOE4 with 10 µM of aspirin during the 16-hour chase period following oleate pulse. We observed that treatment with aspirin during oleate pulse-chase led to an increase in total LD area in APOE3 astrocytes but not APOE4 astrocytes (Fig 4, A-B). This suggests that COX-2 activity at LDs is already reduced in TRAE4 cells, occluding the effect of aspirin, and resulting in accumulation of the COX-2 substrate AA within TGs. We reanalyzed our untargeted lipidomic dataset reported in (Windham et al., 2024), specifically focusing on phospholipid and TG species containing AA. As predicted from the aspirin treatment result, our analysis revealed that TRAE4 astrocytes showed an increase in TGs containing AA, particularly after oleate pulse-chase (Fig. 4, C). This correlates well with our previous finding that PUFA-containing TGs are higher in *APOE4* cells compared to *APOE3* cells (Windham et al., 2024).

**Figure 4.**
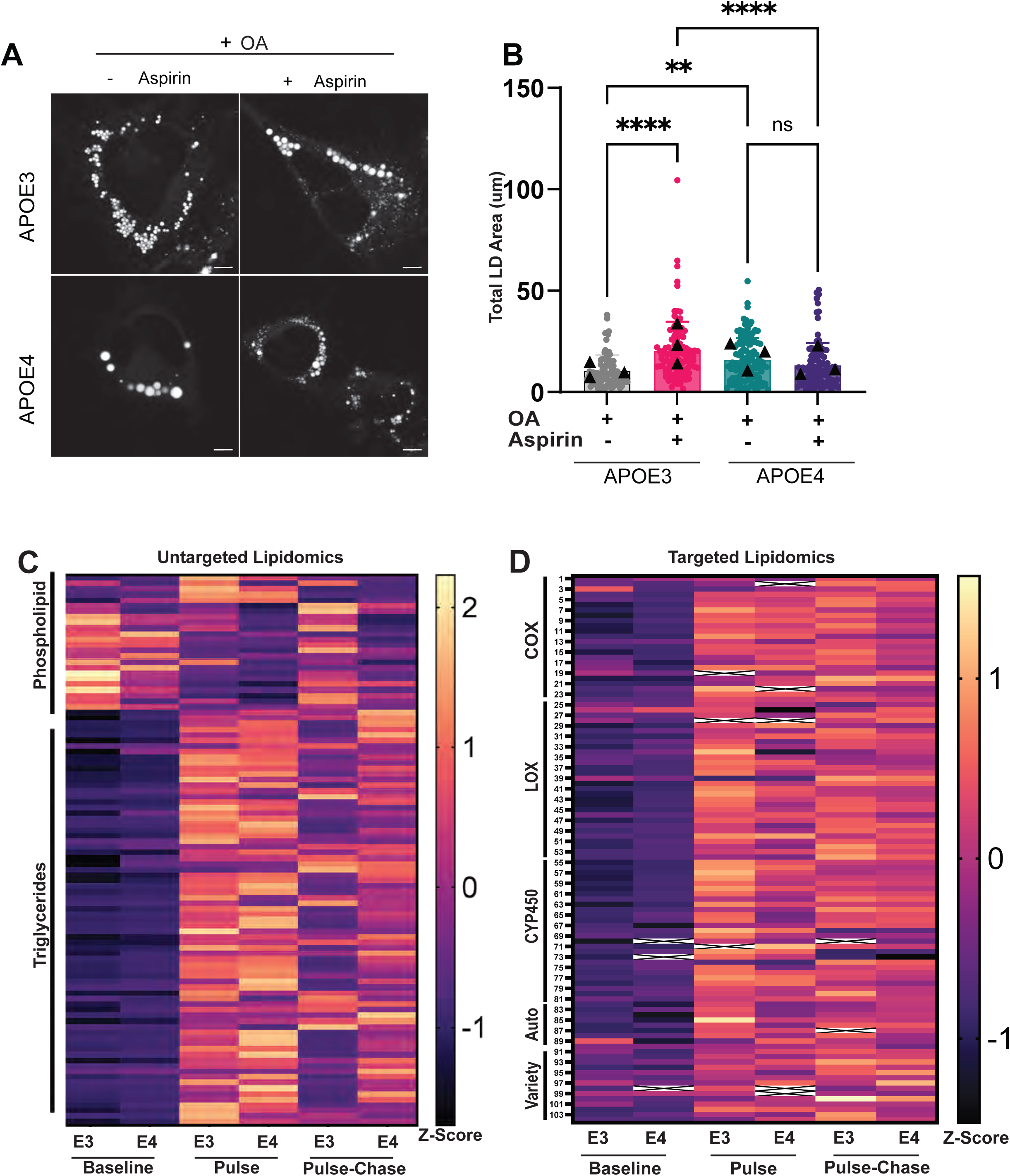
APOE4-expressing cells phenocopy aspirin treatment and exhibit reduced COX-2 activity. (A) Single plane confocal micrographs of live TRAE3 or TRAE4 astrocytes stained for LDs (BODIPY493), either vehicle treated (left) or aspirin treated (right) for 16 hours in media without additional oleate, following a 5 hour 400 µM oleate pulse. (B) Quantification of Total LD area by ImageJ software. Scale bars: 10µm. Groups were compared by one-way ANOVA with a post-hoc Tukey’s t test in Graphpad Prism. (C) Heatmap of arachidonic acid containing phospholipids and triglycerides in TRAE3 or TRAE4 cells at baseline, after oleate pulse, or after oleate pulse-chase, detected by untargeted lipidomics. (D) Heatmap of downstream lipids of arachidonic acid metabolism grouped by predominant producing enzyme in TRAE3 or TRAE4 cells at baseline, after oleate pulse, and after oleate pulse-chase, detected by targeted lipidomics. Individual lipid species are representative to the left of the heatmap as a number, see Table S2. Triangle: Mean of independent biological replicate. Circle: Individual cell measurement across biological replicates. p ≤0.05*, p ≤0.005**, p ≤0.0005***, p ≤0.0001****

### APOE4 blunts the production of intracellular prostaglandins

We hypothesized that the increase in AA-TG species would correlate with reduced production of AA-derived metabolites. However, our untargeted analysis was unable to detect these species. Therefore, to measure AA metabolites, we performed targeted whole cell lipidomic analysis on TRAE3 versus TRAE4 astrocytes at baseline, after oleate pulse, and after oleate pulse-chase. In TRAE3 astrocytes, there was a dramatic increase in most AA metabolites upon oleate pulse, which gradually declined upon removal of excess fatty acid (pulse-chase). Interestingly, we observed a broad reduction in intracellular AA metabolite species in TRAE4 astrocytes following oleate pulse, that appeared to be sustained throughout the pulse-chase (Fig. 4, D, Supplemental Table 2).

We further interrogated the targeted lipidomics data set by quantifying lipid species downstream of specific enzymes. The AA pathway plays a key role in inflammation because it is the precursor for bioactive lipids known as eicosanoids (Wang et al., 2021). Phospholipase A2 releases AA from membranes by hydrolyzing AA esterified in phospholipids. The free AA can then proceed down two predominant metabolic pathways. The first pathway is the cyclooxygenase (COX) pathway, which produces prostanoids (Fig. 5, A). The prostanoid pathway can be further subclassed into prostaglandins, prostacyclins, and thromboxanes. The second pathway that can metabolize free AA is the lipoxygenase (LOX) pathway (Fig. 5, B). The LOX pathway produces subclasses of bioactive lipids, namely lipoxins and leukotrienes (Wang et al., 2021). We observed a marked increase in two subclasses of prostanoids in TRAE3 but not TRAE4 cells upon oleate pulse: prostaglandins and thromboxane (Fig. 5, C-F). To confirm this was specific to the COX pathway and not a general modification of AA metabolism, we also measured lipoxins from the LOX pathway. We observed a significant increase in the production of lipoxins following oleate treatment in both TRAE3 and TRAE4 cells (Fig. 5, G-J), suggesting the reduced production of prostanoids is specific to the APOE-COX-2 interaction. Taken together, our results indicate that APOE4 reduces COX-2 levels at the LD, which leads to reduced intracellular AA metabolites and an accumulation of AA-TGs, effectively sensitizing the astrocytes to lipid peroxidation (Windham et al., 2024; Lange et al., 2021).

**Figure 5.**
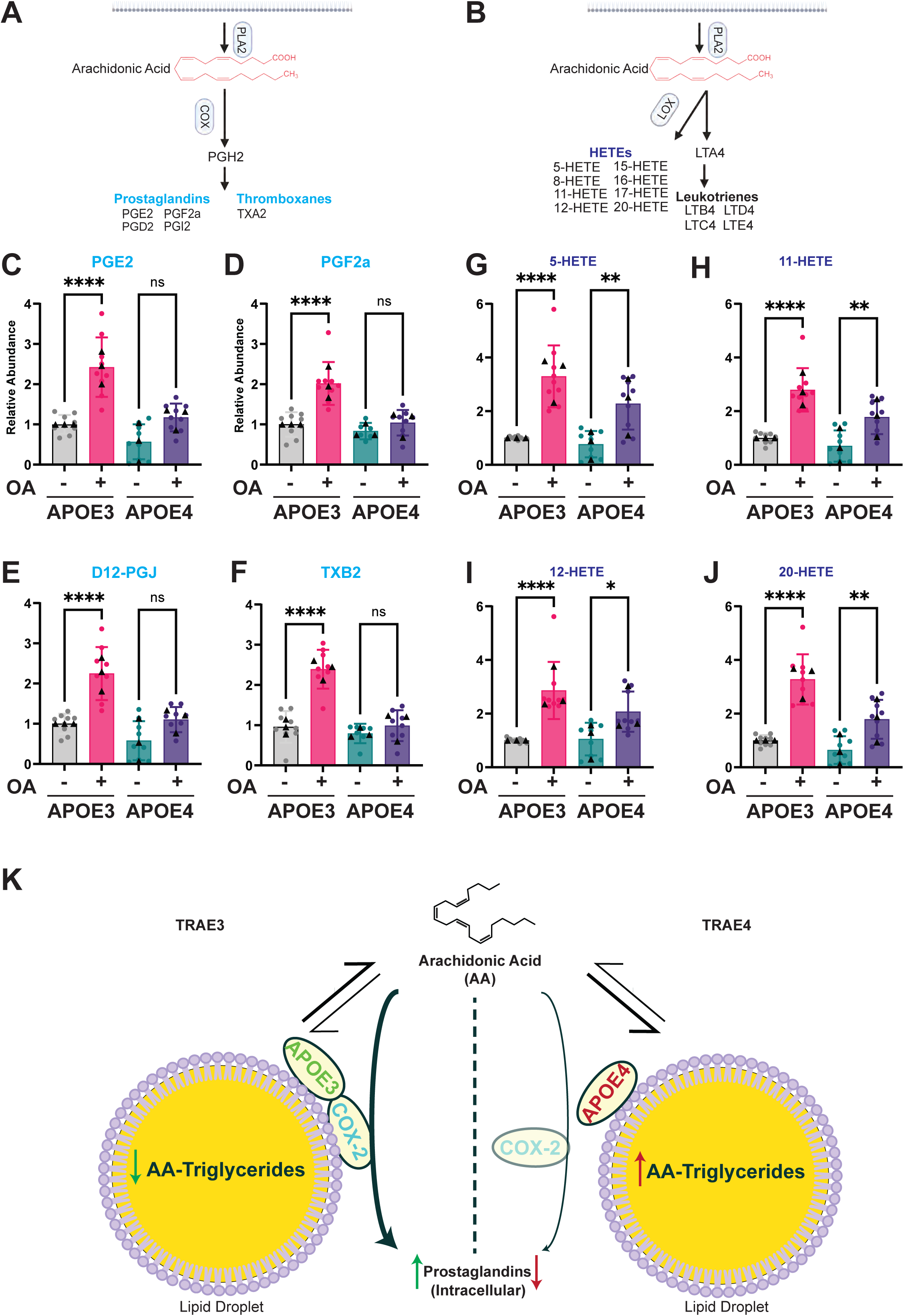
APOE4 expression blunts prostaglandin production downstream of COX-2. (A and B) Schematics of major arachidonic acid metabolite pathways. (C-F) Relative abundance of major prostaglandin species or arachidonic acid direct metabolites downstream of COX-2 in TRAE3 or TRAE4 cells at baseline or after 400 µM oleate pulse, quantified by LC-MS/MS. (G-J) Relative abundance of common lipoxygenase metabolites quantified by LC-MS/MS. (K) Summary schematic of influence of APOE variant expression on COX-2 localization, prostaglandin synthesis, and LD AA-TG accumulation. Triangle: Mean of relative lipid abundance to E3 baseline of independent biological replicates. Circle: Individual relative lipid abundance to E3 baseline measured across biological replicates. Groups were compared by one-way ANOVA with a post-hoc Tukey’s t test in Graphpad Prism. p ≤0.05*, p ≤0.005**, p ≤0.0005***, p ≤0.0001****

## Discussion

We discovered that in states of high lipid load, APOE and COX-2 reroute from the secretory pathway to LDs. In the context of AD, this likely occurs in response to cellular exposure to amyloid beta, which has been shown to induce LD biogenesis in glial cells including astrocytes and microglia, in vitro and in vivo (Liu et al., 2023; Haney et al., 2024; Prakash et al., 2025; Stephens and Johnson, 2025). APOE3 and APOE4 are predicted to bind in distinct orientations with COX-2, with APOE3 promoting the recruitment and/or retention of COX-2 to LDs relative to APOE4. This differential binding and recruitment of COX-2 to the LD between APOE3 versus APOE4 modulates the production and function of prostaglandins, with APOE3 promoting while APOE4 suppresses COX-2 activity at the LD. The LD is the preferred platform to host this APOE-COX-2 driven molecular rewiring, as it concentrates the natural substrate of COX-2, AA, as a TG. The suppression of COX-2 activity by APOE4 has two potentially negative consequences: it reduces the synthesis of intracellular prostaglandins and shunts the AA that accumulates as a result into TGs within LDs, increasing the sensitivity of LDs to lipid peroxidation (Fig. 5, J, Windham et al., 2024).

This modulation of intracellular prostaglandin signaling through physical interaction with COX-2 is likely upstream of many cell-autonomous effects of APOE4 that have been observed in glial cells. Intracellular prostaglandin signaling promotes astrocytes to increase their phagocytic capacities, shifts their intracellular calcium signaling, and alters NF-Kβ expression, effectively priming the astrocyte for reactivity (Dócs et al., 2024). Additionally, prostaglandin receptors have been identified at the nuclear membrane in a variety of cell types where they appear to have distinct molecular functions compared to plasma membrane-localized receptors (Bhattacharya et al., 1998; Fernández-Martínez and Lucio-Cazaña, 2014). Of particular interest is the binding of prostaglandins, particularly PGJ, to PPARs inside the nucleus. This binding is proposed to shift the cell to a state of increased lipid metabolism, reduced pro-inflammatory signaling, and decreased cytokine secretion (Rotondo and Davidson, 2002; Korbecki et al., 2019; Li et al., 2019). Furthermore, both PGA2 and 15-PGJ2, downstream metabolites of PGE2 and PGD2, undergo Michael addition to free thiol groups in cysteines (Musiek et al., 2005). Both IKK-β (an activator of NF-Kβ) and NF-Kβ itself contain cysteines in their active sites (Byun et al., 2006; Perkins, 2012), which are subject to this addition. Addition of PGA2 or 15-PGJ2 to either of these proteins can render them inactive, preventing translocation of NF-Kβ into nucleus, effectively suppressing inflammatory signaling (Straus et al., 2000). Together, these distinctly intracellular actions of prostaglandins have the potential to reprogram astrocytes to reduce tissue wide inflammatory signaling. Consistent with lower intracellular prostaglandin signaling in TRAE4 cells, expression of APOE4 was found to impair immune reactivity in induced pluripotent stem cell (iPSC)-derived astrocytes (Feringa et al., 2025).

In addition to affecting cell-autonomous phenotypes in glia, the localization of APOE and COX-2 to LDs may have implications for non-autonomous cell signaling. Localization of COX-2 to LDs has been suggested to shift prostaglandins to act intracellularly rather than as a secreted lipid signal, switching signaling to an intracrine rather than autocrine/paracrine mode (Horton et al., 1999; Morita et al., 1995; Prescott, 2000; Hartal-Benishay et al., 2024). Therefore, we speculate that the relocalization of APOE and COX-2 to LDs and resulting increase in intracellular prostaglandins in TRAE3 may correlate with decreased secretion of prostaglandins, while TRAE4 astrocytes fail to make this switch and thus secrete higher levels of prostaglandins. Secreted prostaglandins coordinate a tissue-wide amplification of neuroinflammatory signals (Mohri et al., 2006). Thus, TRAE4 astrocytes may become trapped in a paracrine neuroinflammatory signaling cascade that is detrimental to neighboring neurons and glia but fails to properly deal with pathology (Zhang et al., 2025; Guttenplan et al., 2021). This shift has functional consequences on both neighboring neurons and microglia. In myeloid-derived cells, a positive feedback loop between COX2-PGE2 and PGE2 receptors (EP2/4) promotes an autocrine based positive feedback loop that perpetuates a proinflammatory phenotype; activation of these GPCRs increases the production cAMP, activating the CREB pathway, which in turn promotes sustained COX-2 expression and PGE2 secretion (Obermajer and Kalinski, 2012). We speculate that TRAE4 astrocytes participate in a similar feedback loop to amplify neuroinflammatory signaling. Furthermore, PGE2 secretion upregulates the expression of BACE1, a key enzyme in amyloid-beta production, through this same pathway (Gabr et al., 2017). We suggest that this positive feedback loop may also occur in TRAE4 astrocytes, promoting persistent neuroinflammatory and amyloid-beta producing phenotypes. Finally, PGE2 binding to EP2 expressed on microglia impairs phagocytic activity, resulting in increased amyloid-beta accumulation (Shie et al., 2006). This correlates well with recent papers demonstrating expression of APOE4 results in reduced phagocytic capacity, reduced MHC-1 presentation, decreased presentation of the immunoproteasome, and dampened immune responses in glial cells (Feringa et al., 2025; Prakash et al., 2025; Stephenson et al., 2025; Mi et al., 2023; Haney et al., 2024; Liu et al., 2023).

In summary, we discovered that APOE physically interacts with lipid signaling enzyme COX-2 on the surface of LDs. APOE3 promotes, while APOE4 reduces, intracellular prostaglandin production. This interaction reveals a mechanism through which APOE modulates inflammatory lipid signaling in astrocytes, with implications for astrocyte reactivity and function. The APOE-COX-2 interaction may also affect prostaglandin secretion by astrocytes, which in turn could affect microglial and neuronal function. Understanding the mechanism by which APOE modulates inflammatory lipid signaling could lead to the identification of new drug targets for the prevention and treatment of AD.

## Materials and Methods

### Plasmids

mEmerald-N1 (53976; Addgene) was a kind gift from Dr. Michael Davidson (Florida State University, Tallahassee, FL, USA), and pTK881 (219730; Addgene) was a kind gift from Dr. Tal Kafri (UNC-Chapel Hill, Chapel Hill, NC, USA).

APOE3-mEmerald and APOE4-mEmerald were generated for this study. Plasmids containing APOE3 or APOE4 fused to TurboGFP were purchased from Origene (Cat# RG200395 and RG237409). The ORF of APOE was amplified and subcloned into mEmerald-N1 backbone by Gibson assembly using HiFi DNA assembly master mix (E2621, New England Biolabs) to generate APOE3-mEmerald or APOE4-mEmerald. For lentiviral generation, APOE3-mEmerald or APOE4-mEmerald was amplified and subcloned into a pTK881 by Gibson assembly.

### Antibodies and chemicals

The following antibodies were used in this study:

Primary antibodies: recombinant rabbit anti-APOE mAb (1:300 IF; 1:500 WB; Cat# ab52607; Abcam), mouse anti-Tubulin, chicken anti-GFP (Invitrogen, A10262), rabbit anti-cyclooxygenase-2 (1:10,000 WB; 12282S Cell Signaling Technology) and mouse anti-cycloxygenase-2 (1:100 IF; AB300668, Abcam).

Secondary antibodies: donkey anti-rabbit IgG (H+L) Alexa Fluor 568 (1:500 IF; Cat# A10042, RRID:AB_2534017; Thermo Fisher Scientific), donkey anti-mouse IgG (H+L) Alexa Fluor Plus 647 (1:500 IF; Cat# A32787, RRID:AB_2762830; Thermo Fisher Scientific), donkey anti-rabbit IgG (H+L) IRDye 680RD (1:15,000 WB; Cat# 926-68073, RRID:AB_10954442; LI-COR Biosciences), donkey anti-mouse IgG (H+L) (H+L) IRDye 800CW (1:15,000 WB; Cat# 926-32212, RRID:AB_621847; LI-COR Biosciences).

The following chemicals were used in this work: BODIPY 493/503 (Cat# D3922; Thermo Fisher Scientific), BODIPY 665/676 (Cat# B3932; Thermo Fisher Scientific), normal donkey serum (Cat# S30; Sigma-Aldrich), 16% PFA solution (EM grade) (Cat# 15710; EMS), polybrene (Cat# TR-1003-G; Sigma-Aldrich), poly-D lysine (1.0 mg/ml) (Cat# A-003-E; Sigma-Aldrich), blasticidin protease inhibitor cocktail (Cat# P8340; Sigma-Aldrich), proteinase K (Cat# P8107S; NEB), saponin (Cat# AAA1882014; Fisher), and sodium oleate (Cat# O7501; Sigma-Aldrich).

### Cell culture

Immortalized targeted replacement astrocyte (TRAE3-H and TRAE4-H) cell lines were a gift from Dr. Patrick Sullivan of Duke University (described in Morikawa et al., 2005). Targeted replacement astrocytes were maintained in complete medium consisting of Dulbecco’s modified Eagle medium (DMEM) high glucose (Cat# 15-013-CV; Corning), supplemented with 10% fetal bovine serum (FBS, Cat# 97068-085; VWR), 2 mM glutamine (Cat# 25005-CI; Corning), 200 µg/ml Geneticin (Cat# 10131027; Gibco) and 1X penicillin/streptomycin (Cat# 30-002-CI; Corning). All lines were maintained at 37°C and 5% CO_2_.

### Cell transfection

siRNAs were transfected using DharmaFECT 1 (Cat# T-2001; Horizon Discovery) according to the manufacturer’s instructions. 5 µM siRNA was used for each transfection. Non-targeting (NT) siRNA (Cat# D-001810-04), and *APOE* siRNA (Cat# D-006470-04) were purchased from Horizon Discovery. After transfection, cells were incubated in complete medium lacking phenol red and antibiotics for 48 hours (imaging media).

### Transduction of lentiviral APOE-mEmerald constructs

All lentiviral particles were provided by the UNC NeuroTools and Lenti-shRNA Core Facilities. Immortalized astrocytes expressing human APOE (TRAE) were treated with a 1:5 – 1:10 dilution of either APOE3-mEmerald, APOE4-mEmerald, or mEmerald lentiviral particles with 8ug/mL polybrene. After 48 hours, transduced cultures were treated with 12.5ug/mL of blasticidin for 24 hours to select for cells properly expressing the constructs.

### Isolation of APOE-mEmerald complexes for mass spectrometry proteomics

mEmerald protein complex isolation mass spectrometry was performed as previously described, (Miner et al., 2023) utilizing GFP-Trap magnetic beads (Chromotek). Briefly, TRAE3 or TRAE4 expressing APOE-mEmerald constructs (APOE3-mEmerald, APOE4-mEmerald) were washed twice with cold PBS (Thermo Fisher, #14190144), harvested by scraping with a cell lifter, and centrifuged at 350 g for 10 min at 4 ⁰C. Cell pellets were resuspended in 1 mL resuspension buffer (20mM HEPES, pH 7.4, 1.2% polyvinylpyrrolidone) with protease (MiliporeSigma, #8340) and phosphatase inhibitors (MiliporeSigma, #5726 and #P0044), and snap frozen in liquid nitrogen. Cells were lysed by cryogenic grinding using a MM 301 Mixer Mill (10 cycles, 2.5 min at 30Hz) (Retsch). Lysate was resuspended in MS lysis buffer (20 mM K-HEPES pH 7.4, 150 mM NaCl, 100 mM KOAc, 2 mM MgCl2, 0.1% Tween-20, 1 mm ZnCl2 1 mm CaCl2, 0.5% Triton X-100) with protease and phosphatase inhibitors (5 ml/g cells). Resuspended lysate was homogenized using a Polytron (Kinematica) 2 x 15 sec at 25,000 RPM) and pelleted at 2500 g at 4 ⁰C. Cleared lysate was incubated with 50 ul equilibrated GFP-Trap magnetic beads by rotating for 1 h at 4 ⁰C. Beads were washed 6 times with 1 ml of lysis buffer. GFP complexes were eluted from beads in 40 ml 1X LDS Sample Buffer (Invitrogen) at 70 ⁰C for 15 min. Eluted complexes were alkylated with 100 mM iodoacetamide for 1 h at room temperature prior to mass spectrometry analysis.

### Proteomics sample preparation

AP-MS was performed on two biological replicates using different instruments. Immunoprecipitated samples were resolved by SDS-PAGE and visualized by Coomassie staining. Approximately 1 cm gel lanes were excised and destained with 50 mM ammonium bicarbonate:acetonitrile (1:1, v/v), exchanging solution two times every 10 min, then incubating overnight at 4 °C. Destain solution was removed and gel pieces were dehydrated twice with acetonitrile, then dried by vacuum centrifugation. Proteins were reduced with 10 mM dithiothreitol at 56 °C for 10 min followed by 20 min at room temperature, and alkylated with 100 mM iodoacetamide in 25 mM ammonium bicarbonate for 10 min at room temperature in the dark. Gel pieces were washed twice with acetonitrile and dried. Gel pieces were rehydrated in trypsin (20 ng/µL) and digested overnight at 37 °C. Peptides were extracted sequentially with acetonitrile three times, pooled, and dried by vacuum centrifugation. Samples were desalted using C18 spin columns (Pierce) according to the manufacturer’s instructions and dried by vacuum centrifugation, then stored at −80 °C until analysis.

### LC-MS/MS

The peptide samples from replicate 1 (labeled 1082) in Supplemental Table 1 were analyzed by LC-MS/MS using an Easy nLC 1200 coupled to a QExactive HF mass spectrometer equipped with an Easy-Spray Flex source (Thermo Scientific). Samples were injected onto an Easy Spray PepMap C18 column (75 μm id × 25 cm, 2 μm particle size; Thermo Scientific). Peptides were separated over a 65-minute gradient at a 250 nl/min flow rate, consisting of a 50-minute gradient from 5-40% mobile phase B (MPB), and a constant 100% MPB for 15-minutes. Mobile phase A was 0.1% formic acid in water, and mobile phase B consisted of 0.1% formic acid in 80% ACN. The QExactive HF was operated in data-dependent mode, where the 15 most intense precursors were selected for subsequent fragmentation. For full MS scans, m/z was set to 350-1700, resolution set to 120,000, max injection time to 100ms, and AGC target was 3e6. MS/MS scans were acquired using an m/z range of 200 – 2000 m/z, a max IT of 100ms, an isolation window of 1.6mz and a minimum AGC target of 8e3. The normalized collision energy was set to 27% for HCD. Peptide match was set to preferred, and precursors with unknown charge or a charge state of 1 and ≥ 7 were excluded.

The samples from replicate 2 (labeled 1578) in Supplemental Table 1 were injected onto an IonOpticks Aurora series 3 C18 column (75 μm id × 15 cm, 1.6 μm particle size; IonOpticks) and separated over a 30 min method. The gradient for separation consisted of 2-30% mobile phase B at a 300 nl/min flow rate, where mobile phase A was 0.1% formic acid in water and mobile phase B consisted of 0.1% formic acid in 80% ACN. The Orbitrap Astral was operated in Data Dependent Acquisition (DDA) mode, in which the most intense precursors were selected within a 0.6s cycle for subsequent HCD fragmentation and Astral MS/MS detection. For full MS scans, m/z was set to 375 – 1500, resolution was set to 180,000, maximum injection time was 5 ms, and AGC target was 300%. MS/MS scans were acquired in the Astral analyzer using an m/z range of 110 – 2000, a maximum injection time of 2.5 ms, and an AGC target of 100%. The normalized collision energy was set to 30% for HCD, with an isolation window of 2 m/z. Precursors with unknown charge or a charge state of 1 and ≥ 7 were excluded.

### Proteomic data analysis

Raw data files from the QE-HF were processed using MaxQuant v1.6.15.0 and searched against the Uniprot reviewed mouse database (containing 17,051 sequences), appended to a contaminants database, using Andromeda within MaxQuant. Enzyme specificity was set to trypsin, up to two missed cleavage sites were allowed, and methionine oxidation and N-terminus acetylation were set as variable modifications. A 1% FDR was used to filter all data. Match between runs was enabled (5 min match time window, 20 min alignment window), and a minimum of two unique peptides was required for label-free quantitation using the LFQ intensities.

The protein groups file from MaxQuant was imported into Perseus (1.6.14) for further processing (Tyanova et al., 2016). Only proteins with >1 unique+razor peptide were used for LFQ analysis. Proteins with 50% missing values were removed, and missing values were imputed from a normal distribution within Perseus. Log2 fold change (FC) ratios were calculated using the averaged log2 LFQ intensities, and P-values were calculated using a Student’s t-test for each pairwise comparison. Proteins were identified as APOE interactors if they had a Log2 FC ratio greater than 1.0.

Raw data files from the Astral were processed with Fragpipe (v 23.0) using MSFragger (v 4.3) and IonQuant (1.11.11). The files were searched against the mouse SwissProt database downloaded from Uniprot (containing 17,230 entries, downloaded January 2025), appended to the Hao Lab cell culture contaminants database (Frankenfield et al., 2022). Within Fragpipe, the LFQ-MBR workflow was selected, enzyme specificity was set to “stricttrypsin”, up to two missed cleavages were allowed, the minimum peptide length was set to 7, cysteine carbamidomethylation was set as a fixed modification, methionine oxidation, N-terminal acetylation were set as variable modifications. Both precursor and fragment mass tolerances in MSFragger were set to +/- 20ppm. A false discovery rate (FDR) of 1% was used to filter all data. The combined protein file from fragpipe was imported into Perseus (1.6.14) for further processing. Proteins without two valid values in either the E3_OA or E4_OA conditions were removed, and missing values were imputed from a normal distribution within Perseus. Log2 fold change (FC) ratios were calculated using the averaged log2 LFQ intensities, and P-values were calculated using a Student’s t-test for each pairwise comparison.

### Isolation of APOE-mEmerald complexes for co-immunoprecipitation/western blot

Isolation of mEmerald protein complexes for co-immunoprecipitation was performed utilizing the same harvesting method used for mass spectrometry. Following cell harvesting, Co-immunoprecipitation was performed utilizing GFP-Trap magnetic beads (Chromotek) following manufacturer protocols with slight modification. Cell pellets were resuspended in 400 μL of Co-IP Lysis Buffer (10 mM Tris/Cl pH 7.5; 150 mM NaCl; 0.5 mM EDTA; 0.5% NP-40) with protease and phosphatase inhibitors. The cell suspension was then incubated on ice for 30 min with pipetting every 10 min and then pelleted at 20,000 g for 10 min at 4 ⁰C. 600 μl of Co-IP Dilution Buffer (10 mM Tris-Cl pH 7.5; 150 mM NaCl; 0.5 mM EDTA) was added to cleared lysate. Diluted lysate was incubated with 50 μl equilibrated GFP-Trap magnetic beads by rotating for 1 h at 4 ⁰C. Beads were washed 3 times with 500 ul of Co-IP Dilution Buffer. GFP complexes were eluted from beads in 30 μl 6X Laemmli Sample Buffer at 95 ⁰C for 10 min. Protein complexes were probed by Western blotting.

### Immunofluorescence sample preparation

Immunofluorescence was performed as described in Windham et al 2024. In brief, cells were washed twice in 1× PHEM buffer (60 mM PIPES, 27.3 mM HEPES, 8.22 mM MgSO_4_, 10 mM EGTA, and pH 7.0) and fixed for 10 min at room temperature in room temperature–equilibrated 4% PFA in 1× PHEM. After fixation, cells were washed three times in 1× PHEM and then permeabilized in 0.01% saponin in PHEM for 10 min at room temperature. Permeabilization buffer was then replaced with a blocking buffer (10% Normal Donkey Serum, 3% bovine serum albumin [BSA], 300 mM glycine, 0.01% saponin in PHEM) in which cells were incubated 45 min – 1hr at room temperature. The blocking solution was removed, and cells were washed twice with 0.01% saponin in PHEM before applying the primary antibody solution. Primary antibodies were diluted in antibody dilution solution (3% BSA, 0.01% saponin in PHEM). Cells were incubated with primary antibody for 24–48 h at 4°C. After primary antibody incubation, cells were washed with 0.01% saponin three times for 10 min each at room temperature. Secondary antibodies were diluted in antibody dilution solution together with 1 µg/ml BODIPY 493/503 to stain for LDs. Cells were incubated with secondary antibody for 1 h at room temperature and then washed three times in PHEM for 10 min each at room temperature. Cells were imaged immediately after preparing or stored at 4°C and imaged within a week of preparation.

### APOE3/APOE4 conformational ensemble generation

To capture the dynamic structural differences between the APOE3 and APOE4 isoforms (based on the canonical UniProt sequence, P02649), conformational ensembles were generated using BioEmu, a generative deep learning biomolecular emulator trained on over 200 milliseconds of aggregate molecular dynamics (MD) simulation data, structural databases, and experimental protein stability measurements (Lewis et al., 2025). BioEmu employs a diffusion-based generative architecture built upon the AlphaFold2 Evoformer module(Jumper et al., 2021) and produces equilibrium-representative structural ensembles with relative free energy errors of approximately 1 kcal/mol, as benchmarked against millisecond-timescale MD simulation data. The full-length sequences of human APOE3 (C112/R158) and APOE4 (R112/R158) were submitted independently to BioEmu. For each isoform, a large ensemble of backbone conformations (approx. 2000) was generated to comprehensively sample local and global structural flexibility. The C112R substitution defining APOE4 was explicitly encoded in the input sequence to permit BioEmu to capture mutation-induced conformational rearrangements of the four-helix bundle N-terminal domain and its domain–domain interactions with the C-terminal region.

The resulting structural ensembles were analyzed using principal component analysis (PCA) applied to Cα atomic coordinates to reduce the high-dimensional conformational space to its dominant degrees of freedom. The first two principal components (PC1 and PC2), which together describe the largest-variance collective motions of each ensemble, were used as reaction coordinates for visualization of the conformational landscape. Two-dimensional free energy landscapes (FELs) were constructed for both APOE3 and APOE4 ensembles by converting probability densities—estimated via Gaussian kernel density estimation (KDE)—to relative free energies according to ΔG = −*k*B*T* ln(ρ/ρ_max_), where ρ is the population density at each point in the PC1–PC2 space. Conformational clustering was performed using RMSD-based hierarchical clustering of all ensemble members. The centroid structure extracted from the most densely populated global free energy minimum was designated as the representative conformation for each isoform and subsequently used for downstream protein-protein docking.

### Protein–protein docking

Data-driven macromolecular docking of APOE variants against COX-2 homodimer (Lucido et al., 2016,PDB: 5F19) was performed using HADDOCK (High Ambiguity Driven biomolecular DOCKing), version 2.4 (Dominguez et al., 2003). HADDOCK implements an integrative modeling approach in which biochemical and biophysical information is encoded as Ambiguous Interaction Restraints to guide the docking search, followed by simulated annealing in torsion angle space, explicit solvent refinement, and final scoring by a weighted combination of van der Waals, electrostatic, and empirical desolvation energy terms. A 2:1 stoichiometry was modeled, comprising two APOE protomers bound to one COX-2 homodimer, consistent with the known dimeric architecture of cyclooxygenase-2 and the established propensity of APOE to form homodimers in solution. The representative centroid structures of APOE3 and APOE4 derived from their respective BioEmu free energy minima were docked against the human COX-2 homodimer. Docked complex models were grouped by interface RMSD-based clustering, and the top-scoring cluster representatives were selected for downstream structural and electrostatic analysis. The HADDOCK score, which integrates van der Waals interaction energy (E_vdW_), electrostatic energy (E_elec_), and desolvation energy (E_desolv_), was used as the primary ranking criterion.

### Protein–protein interaction analysis

Detailed two-dimensional protein–protein interaction maps for the APOE3–COX-2 and APOE4–COX-2 docked complexes were generated using PDBsum (Laskowski et al., 2018). PDBsum generates schematic diagrams that enumerate and categorize all intermolecular contacts across the binding interface, including hydrogen bonds, electrostatic salt bridges, and non-bonded van der Waals contacts, and identifies the specific donor and acceptor residue pairs involved in each interaction.

The electrostatic potential surface of the COX-2 homodimer was calculated using the Adaptive Poisson-Boltzmann Solver (APBS)(Baker et al., 2001), following protonation state assignment with PDB2PQR using the AMBER force field charge parameters at physiological pH (7.4). To contextualize the docked complexes in their membrane environment, the lipid bilayer boundary was approximated using a layer of spherical pseudo-atoms placed at the predicted membrane insertion depth of COX-2 by the OPM webserver (https://opm.phar.umich.edu/ppm_server3_cgopm)(Lomize et al., 2022). All structural figures were rendered using Chimera-X(Pettersen et al., 2021).

### OA pulse and pulse-chase assays

Oleic Acid (OA) pulse-chase assays were performed as described in Windham et al 2024. For oleic acid pulse experiments cells were treated with 400µM OA or sodium oleate (control) for 5hrs. For pulse-chase experiments OA was removed following the 5hrs treatment and cells were incubated for an additional 16-18hrs in complete medium lacking OA supplementation. Pulse-Chase experiments were additionally treated with either DMSO (0.01%) control or 10µM aspirin during the 16–18-hours media replacement. For *APOE* knockdown COX-2 recruitment assays, 6,000 TRAE3 and TRAE4 cells were seeded in each well. 24 h after seeding, cells were transfected with 5 µM of the indicated siRNA with Dharmafect 1 using the manufacturer’s protocol in imaging media. 48 h after transfection, cells were fixed in 4% PFA in PHEM for 10 minutes at RT. Fixed cells were prepared for immunofluorescence as described above. All conditions were stained for BODIPY, APOE, and COX-2.

For proteomics and lipidomics experiments, cells were seeded in 15-cm cell culture dishes and harvested as described in respective sections at 80-90% confluency, either at baseline or 5 hours after oleate addition.

### Lipidomics sample collection and analysis

TRAE3 or TRAE4 were grown to 60% confluency in 3 x 10-cm cell culture dishes before either receiving vehicle treatment (baseline), 400µM oleate (pulse). Following treatment cells were washed 2x with ice cold PBS and lifted using a cell scraper. Suspended cultures were spun down at 350g for 10 min at 4 C. Pelleted cells were submitted to Wayne State Lipidomics Core for sample processing and “PUFA Metabolome” lipidomic analysis.

### Light microscopy image acquisition

Confocal images were acquired using an inverted Zeiss 800 single-point scanning confocal microscope equipped with 405, 488, 561, and 647 nm diode lasers, two gallium arsenide phosphide (GaAsp) detectors, and one Airyscan detector. Images were acquired using a Plan-Apo 63×/1.4 NA oil objective lens using ZEN Blue software. All live cell imaging was conducted at 37°C and 5% CO_2_.

Image brightness and contrast were adjusted in ImageJ and/or Adobe Photoshop CS.

### Light microscopy image analysis

As described in Windham et al 2024, LD number and size analysis was performed using a semiautomated pipeline in Fiji (Schindelin et al., 2012). Individual cells were first manually segmented and the area outside of each cell was removed using the “Clear Outside” function. To segment LDs, the BODIPY channel was first passed through a Gaussian filter followed by a Laplacian of Gaussian (LoG) filter. The radius of the LoG filter and the Gaussian blur were heuristically optimized for each image to maximize segmentation accuracy. Autothresholding using the Otsu algorithm was then applied, followed by the binary operations “Open,” “Fill Holes,” and “Watershed.” Then, “Analyze Particles” was used to measure the number and size of segmented LDs, as well as the sum area of all LDs in the cell. For quantification of COX-2 LD rings. Individual cells were traced and BODIPY493-positive LDs were autothresholded and expanded by 1 pixel area Fiji; absolute and relative intensity of COX-2 and APOE signal were measured.

Statistical details for all experiments can be found in figure legends. Statistical analysis among groups was performed using Student’s t test.

## Acknowledgements

We would like to thank Patrick Sullivan (Duke University, Durham, NC, USA) and David Holtzmann (Washington University in St. Louis, St. Louis, MO, USA) for kindly providing the APOE targeted replacement astrocyte lines.

Research reported in this publication was supported by the National Institutes of Health under award numbers T32GM133364 (A.E.P.), R01AG081421 (S.C. and L.A.J.), S10RR027926 and S10OD032292 (Lipidomics Core Facility of Wayne State University), and UNC Cancer Center Core Support Grant P30CA016086 (V.R.C.), as well as by generous support from the WoodNext Foundation.

This research is based in part upon work conducted using the UNC Metabolomics and Proteomics Core Facility, which is supported in part by NCI Center Core Support Grant (2P30CA016086-45) to the UNC Lineberger Comprehensive Cancer Center and Nutrition and Obesity Research Center (P30DK056350).

## Author Contributions

A.E. Powers, I.A. Windham, L.A. Johnson, and S. Cohen conceptualized the project and designed the research. A.E. Powers, L.A. Johnson, and S. Cohen acquired funding. K. Kamble performed aspirin pulse-chase experiments. M.G. Nair performed APOE-COX-2 colocalization immunofluorescence experiments. A.E. Powers, I.A. Windham, and C.E. Prim performed protein extraction and affinity-purification for AP-MS experiments. C.A. Mills, S.P. Lyons and L.E. Herring performed mass spectrometry and data analysis for AP-MS experiments. V.R. Chirasani performed computational modeling of COX-2-APOE interactions. A.E. Powers performed all other experiments in the manuscript. A.E. Powers and G.E. Miner wrote ImageJ scripts and performed image analysis. A.E. Powers and S. Cohen wrote the manuscript. S. Cohen supervised the research.

---

**Supplemental Table 1. Proteins associated with APOE as identified by affinity-purification mass spectrometry.** List of proteins associated with APOE3-mEmerald or APOE4-mEmerald in the presence or absence of 400 µM oleic acid. Proteins were identified as high confidence APOE binding partners if Log2 fold change was >1.0 relative to mEmerald controls. Identified proteins from was generated from DDA analysis. Average log2 fold change was used to combine and rank identified proteins.

**Supplemental Table 2. Targeted lipidomics of TRAE3 and TRAE4 astrocytes.** List of intracellular PUFA metabolites identified by liquid-chromatography mass spectrometry in TRAE3 or TRAE4 astrocytes at baseline, after 5h oleate pulse, or after oleate pulse-chase. Detected lipid amounts were normalized to total protein amount across 3 replicates.

## References

Accioly, M.T., P. Pacheco, C.M. Maya-Monteiro, N. Carrossini, B.K. Robbs, S.S. Oliveira, C. Kaufmann, J.A. Morgado-Diaz, P.T. Bozza, and J.P.B. Viola. 2008. Lipid bodies are reservoirs of cyclooxygenase-2 and sites of prostaglandin-E2 synthesis in colon cancer cells. Cancer Res. 68:1732–1740. doi:10.1158/0008-5472.CAN-07-1999.

Alzheimer Alois. 1907. Uber Eine Eigenartige Erkrankung Der Hirnrinde. Allgemei Ne Zeitschriftfur Psychiastre and Psychsch Gerichtlich Medizine. 64:146–148.

Baker, N.A., D. Sept, S. Joseph, M.J. Holst, and J.A. McCammon. 2001. Electrostatics of nanosystems: Application to microtubules and the ribosome. Proc. Natl. Acad. Sci. U. S. A. 98:10037–10041. doi:10.1073/PNAS.181342398;WEBSITE:WEBSITE:PNAS-SITE;WGROUP:STRING:PUBLICATION.

Belloy, M.E., S.J. Andrews, Y. Le Guen, M. Cuccaro, L.A. Farrer, V. Napolioni, and M.D. Greicius. 2023. APOE genotype and Alzheimer disease risk across age, sex, and population ancestry. JAMA Neurol. 80:1284–1294. doi:10.1001/jamaneurol.2023.3599.

Bersuker, K., C.W.H. Peterson, M. To, S.J. Sahl, V. Savikhin, E.A. Grossman, D.K. Nomura, and J.A. Olzmann. 2018. A Proximity Labeling Strategy Provides Insights into the Composition and Dynamics of Lipid Droplet Proteomes. Dev. Cell. 44:97–112.e7. doi:10.1016/j.devcel.2017.11.020.

Bhattacharya, M., K.G. Peri, G. Almazan, A. Ribeiro-Da-Silva, H. Shichi, Y. Durocher, M. Abramovitz, X. Hou, D.R. Varma, and S. Chemtob. 1998. Nuclear localization of prostaglandin E2 receptors. Proc. Natl. Acad. Sci. U. S. A. 95:15792. doi:10.1073/PNAS.95.26.15792.

Bozza, P.T., J.L. Payne, S.G. Morhamt, R. Langenbacht, O. Smithiest, and P.F. Weller. 1996. Leukocyte lipid body formation and eicosanoid generation: Cyclooxygenase-independent inhibition by aspirin (nonsteroidal antiinflammatory agents/cyclooxygenase knockout/leukotrienes/prostaglandin endoperoxide H synthase/lipoxygenase). Medical Sciences Communicated by Seymour J. Klenbanoff. 93:11091–11096.

Byun, M.S., J. Choi, and D.M. Jue. 2006. Cysteine-179 of IκB kinase β plays a critical role in enzyme activation by promoting phosphorylation of activation loop serines. Exp. Mol. Med. 38:546–552. doi:10.1038/EMM.2006.64;KWRD.

Corder, E.H., A.M. Saunders, N.J. Risch, W.J. Strittmatter, D.E. Schmechel, P.C. Gaskell, J.B. Rimmler, P.A. Locke, P.M. Conneally, K.E. Schmader, G.W. Small, A.D. Roses, J.L. Haines, and M.A. Pericak-Vance. 1994. Protective effect of apolipoproteinE type 2 allele for late onset Alzheimer disease. Nat. Genet. 7:180–184. doi:10.1038/ng0694-180.

Cuní-López, C., J.T. Root, Y. Hao, I. Kowal, N. Blomberg, R. Ghirlando, L.G. Yang, S.J. Koppes-den Hertog, M.R. Cookson, R. van der Kant, M. Giera, Y.A. Qi, and P.S. Narayan. 2025. APOE genotypes differentially remodel the astrocytic lipid droplet-associated proteome to shape lipid droplet dynamics. *bioRxiv*. doi:10.1101/2025.08.19.669163.

Dócs, K., A. Balázs, I. Papp, P. Szücs, and Z. Hegyi. 2024. Reactive spinal glia convert 2-AG to prostaglandins to drive aberrant astroglial calcium signaling. Front. Cell. Neurosci. 18:1382465. doi:10.3389/FNCEL.2024.1382465/FULL.

Dominguez, C., R. Boelens, and A.M.J.J. Bonvin. 2003. HADDOCK: a protein-protein docking approach based on biochemical or biophysical information. J. Am. Chem. Soc. 125:1731–1737. doi:10.1021/JA026939X.

Dong, L.M., C. Wilson, M.R. Wardell, T. Simmons, R.W. Mahley, K.H. Weisgraber, and D.A. Agard. 1994. Human apolipoprotein E: Role of arginine 61 in mediating the lipoprotein preferences of the E3 and E4 isoforms. Journal of Biological Chemistry. 269:22358–22365. doi:10.1016/s0021-9258(17)31797-0.

Faustino, A.F., I.C. Martins, F.A. Carvalho, M.A.R.B. Castanho, S. Maurer-Stroh, and N.C. Santos. 2015. Understanding Dengue Virus Capsid Protein Interaction with Key Biological Targets. Sci. Rep. 5:10592. doi:10.1038/SREP10592.

Feringa, F.M., S.J. Koppes-den Hertog, L.Y. Wang, R.J.E. Derks, I. Kruijff, L. Erlebach, J. Heijneman, R. Miramontes, N. Pömpner, N. Blomberg, D. Olivier-Jimenez, L.E. Johansen, A.J. Cammack, A. Giblin, C.E. Toomey, I.V.L. Rose, H. Yuan, M.E. Ward, A.M. Isaacs, M. Kampmann, D. Kronenberg-Versteeg, T. Lashley, L.M. Thompson, A. Ori, Y. Mohammed, M. Giera, and R. van der Kant. 2025. The Neurolipid Atlas: a lipidomics resource for neurodegenerative diseases. Nature Metabolism 2025 7:10. 7:2142–2164. doi:10.1038/s42255-025-01365-z.

Fernández-Martínez, A.B., and F.J. Lucio-Cazaña. 2014. Transactivation of EGFR by prostaglandin E2 receptors: a nuclear story? Cell. Mol. Life Sci. 72:2187. doi:10.1007/S00018-014-1802-1.

Frankenfield, A.M., J. Ni, M. Ahmed, and L. Hao. 2022. Protein Contaminants Matter: Building Universal Protein Contaminant Libraries for DDA and DIA Proteomics. J. Proteome Res. 21:2104–2113. doi:10.1021/ACS.JPROTEOME.2C00145.

Friday, C.M., I.O. Stephens, C.T. Smith, S. Lee, D. Satish, N.A. Devanney, S. Cohen, J.M. Morganti, S.M. Gordon, and L.A. Johnson. 2025. APOE4 reshapes the lipid droplet proteome and modulates microglial inflammatory responses. Neurobiol. Dis. 212. doi:10.1016/j.nbd.2025.106983.

Gabr, A.A., H.J. Lee, X. Onphachanh, Y.H. Jung, J.S. Kim, C.W. Chae, and H.J. Han. 2017. Ethanol-induced PGE2 up-regulates Aβ production through PKA/CREB signaling pathway. Biochimica et Biophysica Acta (BBA) - Molecular Basis of Disease. 1863:2942–2953. doi:10.1016/J.BBADIS.2017.06.020.

Golden, L.R., D.S. Siano, I.O. Stephens, S.M. MacLean, K. Saito, G.L. Nolt, J.L. Funnell, A. V. Pallerla, S. Lee, C. Smith, J. Chen, H. Zhu, C. Voy, C.M. Whitus, G. Hernandez, B.C. Farmer, K. Pandya, D.O. Cowley, S.L. Macauley, S.M. Gordon, J.M. Morganti, and L.A. Johnson. 2025. APOE4 to APOE2 allelic switching in mice improves Alzheimer’s disease-related metabolic signatures, neuropathology and cognition. Nature Neuroscience 2025 28:12. 28:2461–2475. doi:10.1038/s41593-025-02094-y.

Guttenplan, K.A., M.K. Weigel, P. Prakash, P.R. Wijewardhane, P. Hasel, U. Rufen-Blanchette, A.E. Münch, J.A. Blum, J. Fine, M.C. Neal, K.D. Bruce, A.D. Gitler, G. Chopra, S.A. Liddelow, and B.A. Barres. 2021. Neurotoxic reactive astrocytes induce cell death via saturated lipids. Nature 2021 599:7883. 599:102–107. doi:10.1038/s41586-021-03960-y.

Haney, M.S., R. Pálovics, C.N. Munson, C. Long, P.K. Johansson, O. Yip, W. Dong, E. Rawat, E. West, J.C.M. Schlachetzki, A. Tsai, I.H. Guldner, B.S. Lamichhane, A. Smith, N. Schaum, K. Calcuttawala, A. Shin, Y.H. Wang, C. Wang, N. Koutsodendris, G.E. Serrano, T.G. Beach, E.M. Reiman, C.K. Glass, M. Abu-Remaileh, A. Enejder, Y. Huang, and T. Wyss-Coray. 2024. APOE4/4 is linked to damaging lipid droplets in Alzheimer’s disease microglia. Nature 2024 628:8006. 628:154–161. doi:10.1038/s41586-024-07185-7.

Hartal-Benishay, L.H., S. Tal, A.A. Elkader, O. Ehsainieh, R. Srouji-Eid, T. Lavy, O. Kleifeld, M. Mikl, and L. Barki-Harrington. 2024. Activity-dependent COX-2 proteolysis modulates aerobic respiration and proliferation in a prostaglandin-independent manner. iScience. 27:111403. doi:10.1016/j.isci.2024.111403.

Heneka, M.T., W.M. van der Flier, F. Jessen, J. Hoozemanns, D.R. Thal, D. Boche, F. Brosseron, C. Teunissen, H. Zetterberg, A.H. Jacobs, P. Edison, A. Ramirez, C. Cruchaga, J.C. Lambert, A.R. Laza, J.V. Sanchez-Mut, A. Fischer, S. Castro-Gomez, T.D. Stein, L. Kleineidam, M. Wagner, J.J. Neher, C. Cunningham, S.K. Singhrao, M. Prinz, C.K. Glass, J.C.M. Schlachetzki, O. Butovsky, K. Kleemann, P.L. De Jaeger, H. Scheiblich, G.C. Brown, G. Landreth, M. Moutinho, J. Grutzendler, D. Gomez-Nicola, R.M. McManus, K. Andreasson, C. Ising, D. Karabag, D.J. Baker, S.A. Liddelow, A. Verkhratsky, M. Tansey, A. Monsonego, L. Aigner, G. Dorothée, K.A. Nave, M. Simons, G. Constantin, N. Rosenzweig, A. Pascual, G.C. Petzold, J. Kipnis, C. Venegas, M. Colonna, J. Walter, A.J. Tenner, M.K. O’Banion, J.R. Steinert, D.L. Feinstein, M. Sastre, K. Bhaskar, S. Hong, D.P. Schafer, T. Golde, R.M. Ransohoff, D. Morgan, J. Breitner, R. Mancuso, and S.P. Riechers. 2025. Neuroinflammation in Alzheimer disease. Nat. Rev. Immunol. 25:321–352. doi:10.1038/S41577-024-01104-7.

Henne, W.M., and S. Cohen. 2026. Heterogeneity, dynamics and organelle interactions of lipid droplets. Nat. Rev. Mol. Cell Biol. doi:10.1038/S41580-025-00945-X.

Hickenbottom, S.J., A.R. Kimmel, C. Londos, and J.H. Hurley. 2004. Structure of a lipid droplet protein: The PAT family member TIP47. Structure. 12:1199–1207. doi:10.1016/j.str.2004.04.021.

Horton, J.K., A.S. Williams, Z. Smith-Phillips, R.C. Martin, and G. O’Beirne. 1999. Intracellular Measurement of Prostaglandin E2: Effect of Anti-inflammatory Drugs on Cyclooxygenase Activity and Prostanoid Expression. Anal. Biochem. 271:18–28. doi:10.1006/ABIO.1999.4118.

Ioannou, M.S., J. Jackson, S.H. Sheu, C.L. Chang, A. V. Weigel, H. Liu, H.A. Pasolli, C.S. Xu, S. Pang, D. Matthies, H.F. Hess, J. Lippincott-Schwartz, and Z. Liu. 2019. Neuron-Astrocyte Metabolic Coupling Protects against Activity-Induced Fatty Acid Toxicity. Cell. 177:1522–1535.e14. doi:10.1016/J.CELL.2019.04.001.

Jarc, E., and T. Petan. 2020a. A twist of FATe: Lipid droplets and inflammatory lipid mediators. Biochimie. 169:69–87. doi:10.1016/j.biochi.2019.11.016.

Jarc, E., and T. Petan. 2020b. A twist of FATe: Lipid droplets and inflammatory lipid mediators. Biochimie. 169:69–87. doi:10.1016/j.biochi.2019.11.016.

Jumper, J., R. Evans, A. Pritzel, T. Green, M. Figurnov, O. Ronneberger, K. Tunyasuvunakool, R. Bates, A. Žídek, A. Potapenko, A. Bridgland, C. Meyer, S.A.A. Kohl, A.J. Ballard, A. Cowie, B. Romera-Paredes, S. Nikolov, R. Jain, J. Adler, T. Back, S. Petersen, D. Reiman, E. Clancy, M. Zielinski, M. Steinegger, M. Pacholska, T. Berghammer, S. Bodenstein, D. Silver, O. Vinyals, A.W. Senior, K. Kavukcuoglu, P. Kohli, and D. Hassabis. 2021. Highly accurate protein structure prediction with AlphaFold. Nature. 596:583–589. doi:10.1038/S41586-021-03819-2.

Korbecki, J., R. Bobiński, and M. Dutka. 2019. Self-regulation of the inflammatory response by peroxisome proliferator-activated receptors. Inflammation Research 2019 68:6. 68:443–458. doi:10.1007/S00011-019-01231-1.

Lange, M., P.V. Wagner, and M. Fedorova. 2021. Lipid composition dictates the rate of lipid peroxidation in artificial lipid droplets. Free Radic. Res. 55:469–480. doi:10.1080/10715762.2021.1898603.

Laskowski, R.A., J. Jabłońska, L. Pravda, R.S. Vařeková, and J.M. Thornton. 2018. PDBsum: Structural summaries of PDB entries. Protein Sci. 27:129–134. doi:10.1002/PRO.3289.

Lee, H., R. V. Pearse, A.M. Lish, C. Pan, Z.M. Augur, G. Terzioglu, P. Gaur, M. Liao, M. Fujita, E.S. Tio, D.M. Duong, D. Felsky, N.T. Seyfried, V. Menon, D.A. Bennett, P.L. De Jager, and T.L. Young-Pearse. 2025. Contributions of Genetic Variation in Astrocytes to Cell and Molecular Mechanisms of Risk and Resilience to Late-Onset Alzheimer’s Disease. Glia. 73:1166–1187. doi:10.1002/GLIA.24677.

Lewis, S., T. Hempel, J. Jiménez-Luna, M. Gastegger, Y. Xie, A.Y.K. Foong, V. García Satorras, O. Abdin, B.S. Veeling, I. Zaporozhets, Y. Chen, S. Yang, A.E. Foster, A. Schneuing, J. Nigam, F. Barbero, V. Stimper, A. Campbell, J. Yim, M. Lienen, Y. Shi, S. Zheng, H. Schulz, U. Munir, R. Sordillo, R. Tomioka, C. Clementi, and F. Noé. 2025. Scalable emulation of protein equilibrium ensembles with generative deep learning. Science. 389. doi:10.1126/SCIENCE.ADV9817.

Li, J., C. Guo, and J. Wu. 2019. 15-Deoxy-Δ-12,14-Prostaglandin J2 (15d-PGJ2), an Endogenous Ligand of PPAR-γ: Function and Mechanism. PPAR Res. 2019:7242030. doi:10.1155/2019/7242030.

Liu, C.C., N. Wang, Y. Chen, Y. Inoue, F. Shue, Y. Ren, M. Wang, W. Qiao, T.C. Ikezu, Z. Li, J. Zhao, Y. Martens, S. V. Doss, C.L. Rosenberg, S. Jeevaratnam, L. Jia, A.C. Raulin, F. Qi, Y. Zhu, A. Alnobani, J. Knight, Y. Chen, C. Linares, A. Kurti, J.D. Fryer, B. Zhang, L.J. Wu, B.Y.S. Kim, and G. Bu. 2023. Cell-autonomous effects of APOE4 in restricting microglial response in brain homeostasis and Alzheimer’s disease. Nat. Immunol. 24:1854–1866. doi:10.1038/S41590-023-01640-9.

Liu, L., K. Zhang, H. Sandoval, S. Yamamoto, M. Jaiswal, E. Sanz, Z. Li, J. Hui, B.H. Graham, A. Quintana, and H.J. Bellen. 2015. Glial lipid droplets and ROS induced by mitochondrial defects promote neurodegeneration. Cell. 160:177–190. doi:10.1016/j.cell.2014.12.019.

Lomize, A.L., S.C. Todd, and I.D. Pogozheva. 2022. Spatial arrangement of proteins in planar and curved membranes by PPM 3.0. Protein Sci. 31:209–220. doi:10.1002/PRO.4219.

Lucido, M.J., B.J. Orlando, A.J. Vecchio, and M.G. Malkowski. 2016. Crystal Structure of Aspirin-Acetylated Human Cyclooxygenase-2: Insight into the Formation of Products with Reversed Stereochemistry. Biochemistry. 55:1226–1238. doi:10.1021/ACS.BIOCHEM.5B01378.

MacPherson, R.E.K., S. V. Ramos, R. Vandenboom, B.D. Roy, and S.J. Peters. 2013. Skeletal muscle PLIN proteins, ATGL and CGI-58, interactions at rest and following stimulated contraction. Am. J. Physiol. Regul. Integr. Comp. Physiol. 304:644–650. doi:10.1152/AJPREGU.00418.2012.

Marschallinger, J., T. Iram, M. Zardeneta, S.E. Lee, B. Lehallier, M.S. Haney, J. V. Pluvinage, V. Mathur, O. Hahn, D.W. Morgens, J. Kim, J. Tevini, T.K. Felder, H. Wolinski, C.R. Bertozzi, M.C. Bassik, L. Aigner, and T. Wyss-Coray. 2020. Lipid-droplet-accumulating microglia represent a dysfunctional and proinflammatory state in the aging brain. Nat. Neurosci. 23:194–208. doi:10.1038/S41593-019-0566-1;SUBJMETA.

Mauch, D.H., K. Nägier, S. Schumacher, C. Göritz, E.C. Müller, A. Otto, and F.W. Pfrieger. 2001. CNS synaptogenesis promoted by glia-derived cholesterol. Science. 294:1354–1357. doi:10.1126/SCIENCE.294.5545.1354.

Mi, Y., G. Qi, F. Vitali, Y. Shang, A.C. Raikes, T. Wang, Y. Jin, R.D. Brinton, H. Gu, and F. Yin. 2023. Loss of fatty acid degradation by astrocytic mitochondria triggers neuroinflammation and neurodegeneration. Nat. Metab. 5:445–465. doi:10.1038/S42255-023-00756-4.

Michael Garavito, R., M.G. Malkowski, and D.L. DeWitt. 2002. The structures of prostaglandin endoperoxide H synthases-1 and -2. Prostaglandins Other Lipid Mediat. 68–69:129–152. doi:10.1016/S0090-6980(02)00026-6.

Miner, G.E., C.M. So, W. Edwards, R.A. Coleman, E.L. Klett, J. V Ragusa, J.T. Wine, D. Wong Gutierrez, M. V Airola, L.E. Herring, and S. Cohen. 2023. PLIN5 interacts with FATP4 at membrane contact sites to promote lipid droplet-to-mitochondria fatty acid transport. Dev. Cell. 58:1250–1265.e6. doi:10.1016/j.devcel.2023.05.006.

Mohri, I., M. Taniike, H. Taniguchi, T. Kanekiyo, K. Aritake, T. Inui, N. Fukumoto, N. Eguchi, A. Kushi, H. Sasai, Y. Kanaoka, K. Ozono, S. Narumiya, K. Suzuki, and Y. Urade. 2006. Prostaglandin D2-Mediated Microglia/Astrocyte Interaction Enhances Astrogliosis and Demyelination in twitcher. The Journal of Neuroscience. 26:4383. doi:10.1523/JNEUROSCI.4531-05.2006.

Morikawa, M., J.D. Fryer, P.M. Sullivan, E.A. Christopher, S.E. Wahrle, R.B. DeMattos, M.A. O’Dell, A.M. Fagan, H.A. Lashuel, T. Walz, K. Asai, and D.M. Holtzman. 2005. Production and characterization of astrocyte-derived human apolipoprotein E isoforms from immortalized astrocytes and their interactions with amyloid-β. Neurobiol. Dis. 19:66–76. doi:10.1016/j.nbd.2004.11.005.

Morita, I., M. Schindler, M.K. Regier, J.C. Otto, T. Hori, D.L. DeWitt, and W.L. Smith. 1995. Different Intracellular Locations for Prostaglandin Endoperoxide H Synthase-1 and −2. Journal of Biological Chemistry. 270:10902–10908. doi:10.1074/JBC.270.18.10902.

Musiek, E.S., L. Gao, G.L. Milne, W. Han, M.B. Everhart, D. Wang, M.G. Backlund, R.N. DuBois, G. Zanoni, G. Vidari, T.S. Blackwell, and J.D. Morrow. 2005. Cyclopentenone isoprostanes inhibit the inflammatory response in macrophages. Journal of Biological Chemistry. 280:35562–35570. doi:10.1074/jbc.M504785200.

Nakato, M., M. Matsuo, N. Kono, M. Arita, H. Arai, J. Ogawa, N. Kioka, and K. Ueda. 2015. Neurite outgrowth stimulation by n-3 and n-6 PUFAs of phospholipids in apoE-containing lipoproteins secreted from glial cells. J. Lipid Res. 56:1880–1890. doi:10.1194/jlr.M058164.

Nemergut, M., S.M. Marques, L. Uhrik, T. Vanova, M. Nezvedova, D.C. Gadara, D. Jha, J. Tulis, V. Novakova, J. Planas-Iglesias, A. Kunka, A. Legrand, H. Hribkova, V. Pospisilova, J. Sedmik, J. Raska, Z. Prokop, J. Damborsky, D. Bohaciakova, Z. Spacil, L. Hernychova, D. Bednar, and M. Marek. 2023. Domino-like effect of C112R mutation on ApoE4 aggregation and its reduction by Alzheimer’s Disease drug candidate. Mol. Neurodegener. 18. doi:10.1186/S13024-023-00620-9.

Nguyen, D., P. Dhanasekaran, M. Nickel, C. Mizuguchi, M. Watanabe, H. Saito, M.C. Phillips, and S. Lund-Katz. 2014. Influence of domain stability on the properties of human apolipoprotein E3 and E4 and mouse apolipoprotein E. Biochemistry. 53:4025–4033. doi:10.1021/BI500340Z.

Obermajer, N., and P. Kalinski. 2012. Key role of the positive feedback between PGE2 and COX2 in the biology of myeloid-derived suppressor cells. Oncoimmunology. 1:762–764. doi:10.4161/ONCI.19681.

Olzmann, J.A., and P. Carvalho. 2019. Dynamics and functions of lipid droplets. Nat. Rev. Mol. Cell Biol. 20:137–155. doi:10.1038/s41580-018-0085-z.

Perkins, N.D. 2012. Cysteine 38 holds the key to NF-κB activation. Mol. Cell. 45:1–3. doi:10.1016/j.molcel.2011.12.023.

Pettersen, E.F., T.D. Goddard, C.C. Huang, E.C. Meng, G.S. Couch, T.I. Croll, J.H. Morris, and T.E. Ferrin. 2021. UCSF ChimeraX: Structure visualization for researchers, educators, and developers. Protein Sci. 30:70–82. doi:10.1002/PRO.3943.

Pfrieger, F.W., and B.A. Barres. 1997. Synaptic efficacy enhanced by glial cells in vitro. Science. 277:1684–1687. doi:10.1126/SCIENCE.277.5332.1684.

Prakash, P., P. Manchanda, E. Paouri, K. Bisht, K. Sharma, J. Rajpoot, V. Wendt, A. Hossain, P.R. Wijewardhane, C.E. Randolph, Y. Chen, S. Stanko, N. Gasmi, A. Gjojdeshi, S. Card, J. Fine, K.P. Jethava, M.G. Clark, B. Dong, S. Ma, A. Crockett, E.A. Thayer, M. Nicolas, R. Davis, D. Hardikar, D. Allende, R.A. Prayson, C. Zhang, D. Davalos, and G. Chopra. 2025. Amyloid-β induces lipid droplet-mediated microglial dysfunction via the enzyme DGAT2 in Alzheimer’s disease. Immunity. 58:1536–1552.e8. doi:10.1016/j.immuni.2025.04.029.

Prescott, S.M. 2000. Is cyclooxygenase-2 the alpha and the omega in cancer? J. Clin. Invest. 105:1511–1513. doi:10.1172/JCI10241.

Ralhan, I., C.L. Chang, J. Lippincott-Schwartz, and M.S. Ioannou. 2021. Lipid droplets in the nervous system. J. Cell Biol. 220. doi:10.1083/JCB.202102136.

Ratnavadivel, S., K. Jürgens, N. Klinke, A. Gärtner, J. Groß, K. Klingel, A. Malmendal, S. Walter, H. Boen, E. Van Craenenbroeck, R. Schramm, A. Kostareva, J. Gummert, A. Kassner, H. Meyer, A. Paululat, and H. Milting. 2026. Newfoundland Mutation TMEM43-p.S358L Causes Impaired Cardiac Energy Metabolism and Mitochondrial Function Through Altered Protein Interaction. Circulation: Genomic and Precision Medicine. doi:10.1161/CIRCGEN.125.005171;CTYPE:STRING:JOURNAL.

Raulin, A.C., S. V. Doss, Z.A. Trottier, T.C. Ikezu, G. Bu, and C.C. Liu. 2022. ApoE in Alzheimer’s disease: pathophysiology and therapeutic strategies. Mol. Neurodegener. 17. doi:10.1186/S13024-022-00574-4.

Rotondo, D., and J. Davidson. 2002. Prostaglandin and PPAR control of immune cell function. Immunology. 105:20. doi:10.1046/J.0019-2805.2001.01361.X.

Rueter, J., G. Rimbach, C. Treitz, A. Schloesser, K. Lüersen, A. Tholey, and P. Huebbe. 2023. The mitochondrial BCKD complex interacts with hepatic apolipoprotein E in cultured cells in vitro and mouse livers in vivo. Cellular and Molecular Life Sciences 2023 80:3. 80:59-. doi:10.1007/S00018-023-04706-X.

Schindelin, J., I. Arganda-Carreras, E. Frise, V. Kaynig, M. Longair, T. Pietzsch, S. Preibisch, C. Rueden, S. Saalfeld, B. Schmid, J.Y. Tinevez, D.J. White, V. Hartenstein, K. Eliceiri, P. Tomancak, and A. Cardona. 2012. Fiji: an open-source platform for biological-image analysis. Nat. Methods. 9:676–682. doi:10.1038/NMETH.2019.

Shie, F.S., K.S. Montine, R.M. Breyer, and T.J. Montine. 2006. Microglial EP2 as a New Target to Increase Amyloid β Phagocytosis and Decrease Amyloid β-Induced Damage to Neurons. Brain Pathology. 15:134. doi:10.1111/J.1750-3639.2005.TB00509.X.

Stelzmann, R.A., H. Norman Schnitzlein, and F. Reed Murtagh. 1995. An English translation of Alzheimer’s 1907 paper, “Uber eine eigenartige Erkankung der Hirnrinde.” Clin. Anat. 8:429–431. doi:10.1002/CA.980080612.

Stephens, I.O., and L.A. Johnson. 2025. Knockout of Perilipin-2 in Microglia Alters Lipid Droplet Accumulation and Response to Alzheimer’s Disease Stimuli. Cells. 14:1783. doi:10.3390/CELLS14221783/S1.

Stephenson, R.A., J. Sepulveda, K.R. Johnson, A. Lita, J. Gopalakrishnan, D.J. Acri, A. Beilina, L. Cheng, L.G. Yang, J.T. Root, M.E. Ward, C. Combs, W.C. Skarnes, M.R. Cookson, H.Y. Shih, M. Larion, G.W. Rebeck, and P.S. Narayan. 2025. Triglyceride metabolism controls inflammation and microglial phenotypes associated with APOE4. Cell Rep. 44. doi:10.1016/j.celrep.2025.115961.

Straus, D.S., G. Pascual, M. Li, J.S. Welch, M. Ricote, C.H. Hsiang, L.L. Sengchanthalangsy, G. Ghosh, and C.K. Glass. 2000. 15-Deoxy-Δ12,14-prostaglandin J2 inhibits multiple steps in the NF-κB signaling pathway. Proc. Natl. Acad. Sci. U. S. A. 97:4844–4849. doi:10.1073/PNAS.97.9.4844;SUBPAGE:STRING:FULL.

Thiam, A.R., R. V. Farese, and T.C. Walther. 2013. The biophysics and cell biology of lipid droplets. Nat. Rev. Mol. Cell Biol. 14:775–786. doi:10.1038/nrm3699.

Tyanova, S., T. Temu, P. Sinitcyn, A. Carlson, M.Y. Hein, T. Geiger, M. Mann, and J. Cox. 2016. The Perseus computational platform for comprehensive analysis of (prote)omics data. Nat. Methods. 13:731–740. doi:10.1038/NMETH.3901.

Wang, B., L. Wu, J. Chen, L. Dong, C. Chen, Z. Wen, J. Hu, I. Fleming, and D.W. Wang. 2021. Metabolism pathways of arachidonic acids: mechanisms and potential therapeutic targets. Signal Transduction and Targeted Therapy 2021 6:1. 6:94-. doi:10.1038/s41392-020-00443-w.

Weisgraber$, K.H., S.C. Rall, and R.W. Mahley. 1981. Human E apoprotein heterogeneity. Cysteine-arginine interchanges in the amino acid sequence of the apo-E isoforms. Journal of Biological Chemistry. 256:9077–9083. doi:10.1016/S0021-9258(19)52510-8.

Windham, I.A., and S. Cohen. 2024. The cell biology of APOE in the brain. Trends Cell Biol. 34:338–348. doi:10.1016/j.tcb.2023.09.004.

Windham, I.A., A.E. Powers, J. V. Ragusa, E.D. Wallace, M.C. Zanellati, V.H. Williams, C.H. Wagner, K.K. White, and S. Cohen. 2024. APOE traffics to astrocyte lipid droplets and modulates triglyceride saturation and droplet size. Journal of Cell Biology. 223. doi:10.1083/jcb.202305003.

Wu, X., J.A. Miller, B.T.K. Lee, Y. Wang, and C. Ruedl. 2025. Reducing microglial lipid load enhances β amyloid phagocytosis in an Alzheimer’s disease mouse model. Sci. Adv. 11:6038. doi:10.1126/SCIADV.ADQ6038.

Xu, Q., A. Bernardo, D. Walker, T. Kanegawa, R.W. Mahley, and Y. Huang. 2006a. Profile and regulation of apolipoprotein E (ApoE) expression in the CNS in mice with targeting of green fluorescent protein gene to the ApoE locus. J. Neurosci. 26:4985–4994. doi:10.1523/JNEUROSCI.5476-05.2006.

Xu, Q., A. Bernardo, D. Walker, T. Kanegawa, R.W. Mahley, and Y. Huang. 2006b. Profile and Regulation of Apolipoprotein E (ApoE) Expression in the CNS in Mice with Targeting of Green Fluorescent Protein Gene to the ApoE Locus. Journal of Neuroscience. 26:4985–4994. doi:10.1523/JNEUROSCI.5476-05.2006.

Zhang, Z., K. Yu, H. Bao, T.C. Ikezu, A.R. Ravula, B. Melvin, C. Scholes, Y. You, J. Ellison, T. Kanekiyo, M.A. DeTure, D.W. Dickson, X. Han, J. Peng, S. Ikezu, and T. Ikezu. 2025. APOE4 drives neuroinflammation and lipid dysbiosis in Alzheimer’s disease by modulating lipid compositions and cell adhesion molecules in brain-derived extracellular vesicles. Alzheimer’s & Dementia. 21:e105076. doi:10.1002/ALZ70855_105076.

Zheng, Q., and X. Wang. 2025. Alzheimer’s disease: insights into pathology, molecular mechanisms, and therapy. Protein Cell. 16:83–120. doi:10.1093/procel/pwae026.

